# A conserved immune trajectory of recovery in hospitalized COVID-19 patients

**DOI:** 10.1101/2022.03.15.484467

**Authors:** Cassandra E. Burnett, Trine Line Hauge Okholm, Iliana Tenvooren, Diana M. Marquez, Stanley Tamaki, Priscila Munoz Sandoval, The UCSF COMET Consortium, Carolyn S. Calfee, Carolyn M. Hendrickson, Kirsten N. Kangelaris, Charles R. Langelier, Matthew F. Krummel, Prescott G. Woodruff, David J. Erle, K. Mark Ansel, Matthew H. Spitzer

## Abstract

Many studies have provided insights into the immune response to COVID-19; however, little is known about the immunological changes and immune signaling occurring during COVID-19 resolution. Individual heterogeneity and variable disease resolution timelines obscure unifying immune characteristics. Here, we collected and profiled >200 longitudinal peripheral blood samples from patients hospitalized with COVID-19, with other respiratory infections, and healthy individuals, using mass cytometry to measure immune cells and signaling states at single cell resolution. COVID-19 patients showed a unique immune composition and an early, coordinated and elevated immune cell signaling profile, which correlated with early hospital discharge. Intra-patient time course analysis tied to clinically relevant events of recovery revealed a conserved set of immunological processes that accompany, and are unique to, disease resolution and discharge. This immunological process, together with additional changes in CD4 regulatory T cells and basophils, accompanies recovery from respiratory failure and is associated with better clinical outcomes at the time of admission. Our work elucidates the biological timeline of immune recovery from COVID-19 and provides insights into the fundamental processes of COVID-19 resolution in hospitalized patients.

## Introduction

SARS-CoV-2 and the resulting disease COVID-19 has resulted in over 244,000,000 infected individuals and more than 4,900,000 deaths globally as of October 28th, 2021 (World Health Organization 2021b). Most people infected with SARS-CoV-2 are asymptomatic or experience mild flu-like symptoms. In a prospective study of adults confirmed with SARS-CoV2, 91% of patients were asymptomatic or were outpatients with mild illness, while only 9% required inpatient care (Logue et al. 2021). These patients can develop severe disease, including pneumonia, acute respiratory distress syndrome (ARDS), or multiple organ failure, and often require supplemental oxygen support or, in the most critical cases, mechanical ventilation. Although a small percentage of all infected patients succumb to the disease (1.3%) (Centers for Disease Control and Prevention 2021), the majority of hospitalized patients are able to successfully combat and clear the infection. Many studies have focused on features defining the subset of patients who ultimately succumb to disease, however, it is also essential to profile a successful resolution and identify conserved immune features during this interval to understand the majority of patient responses.

The immunopathology of COVID-19 has broadly been characterized by lymphopenia, lymphocyte dysfunction, abnormalities of innate immune cells, and increased cytokine production (Mathew et al. 2020; Mann et al. 2020; Lucas et al. 2020; Yang et al. 2020). Early observations of serum cytokine levels in COVID-19 patients revealed high levels of circulating IL-6, generating the hypothesis of an IL-6 driven cytokine storm and resulting immunopathology (Moore and June 2020; Yang et al. 2020). Analysis of clinical trials using IL-6 neutralizing therapies demonstrate appreciable clinical benefits with additional benefits observed in patients receiving additional corticosteroid treatment suggesting modulation of immune signaling and immune cell activation has clinical implications for disease escalation and resolution (Rosas et al. 2021; Tsai et al. 2020; WHO Rapid Evidence Appraisal for COVID-19 Therapies (REACT) Working Group et al. 2021). Additionally, insufficient type I interferon (IFN) signaling and autoantibodies that inhibit type I IFN have been linked to a subset of severe cases of COVID-19, suggesting that type I immune responses and IFN signaling are likely protective (Combes et al. 2021; Wang et al. 2021; van der Wijst et al. 2021; Chang et al. 2021; Q. Zhang et al. 2020; Asano et al. 2021). High serum cytokine levels along with observations of broad immunological misfiring have been observed across patient subsets, indicating a delicate balance between productive and destructive immune responses and suggesting the importance of evaluating immune cell signaling. However, it remains unclear what, if any, immune cell signaling is protective and how immune cell signaling dynamics change over time in patients who resolve or fail to resolve COVID-19.

While many studies have made significant contributions to our understanding of the immune system and its relation to COVID-19, most analytical approaches are cross-sectional and describe the immunological differences between COVID-19 severity groups defined by clinical metrics, such as the WHO score. In comparison, longitudinal studies are uniquely capable of assessing changes in the immune response during disease progression or resolution over time. Elucidating the immunological events that accompany successful disease resolution is essential to inform the management of patient care and contextualize the deviations from successful resolution that characterize the most severe disease cases. Because the infection timeline is highly variable, and human immunological responses are diverse, understanding immunological dynamics during this specific recovery period requires longitudinal monitoring and high-dimensional data from a large cohort of patients.

Here, we investigated intra-patient immunological changes across clinically relevant time points to identify changes in immune responses that accompany effective COVID-19 resolution. We obtained longitudinal peripheral blood samples (n = 230) from hospitalized COVID-19 patients, SARS-CoV-2 negative ventilated patients, and healthy individuals. To investigate changes in immune cell signaling states over time, we utilized mass cytometry with a unique panel of antibodies specific for immune cell phenotyping and for measuring phosphorylated cell signaling proteins. We identified distinct immune cell composition and signaling states in COVID-19 patients compared to COVID-19 negative patients and healthy individuals. Additionally, we discovered a conserved and coordinated immune response that accompanies COVID-19 resolution and hospital discharge. Furthermore, these and other features were relevant to resolution of the most severe mechanically ventilated patients, and these immune cell states correlated with better clinical outcomes at time of admission. Our findings indicate that, although patients have heterogeneous immunological baselines and highly variable disease courses, there exists a core immunological trajectory that defines recovery from severe SARS-CoV-2 infection. Our results provide a working model of a successful immune response trajectory among patients with COVID-19 requiring hospitalization, deviations from which are associated with extended hospitalization and mortality.

## Results

### Longitudinal peripheral blood sampling from hospitalized COVID-19 positive and negative patients

To investigate the composition of circulating immune cells and the cell signaling states that characterize SARS-COV-2 infections and distinguish it from other respiratory infections, we collected longitudinal peripheral blood (PB) samples from COVID-19 patients and COVID-19 negative patients (PCR negative for SARS-COV-2) admitted to UCSF Medical Center and Zuckerberg San Francisco General Hospital. PB samples and corresponding patient demographics and clinical parameters, e.g. World Health Organization (WHO) severity scores (World Health Organization 2021a), ventilation duration, and hospital length of stay, were collected throughout inpatient care (Table S1 and S2). PB samples from healthy individuals (n = 11) were obtained as controls (Table S3). All samples were processed, stained, and analyzed by mass cytometry to quantify the expression of 30 protein markers and 14 phosphorylated signaling molecules (Table S4). Samples that met our quality control standards (methods) were normalized across batches to obtain our final cohort of 230 samples; 205 samples from 81 COVID-19 patients, 14 samples from 7 COVID-19 negative patients, and single samples from each of 11 healthy individuals (Figure 1A and S1A). COVID-19 patients were classified into COVID-19 severity groups based on their WHO score at day of sampling (3: mild, 4: moderate, 5-6-7: severe) (World Health Organization 2021a). Based on the phenotypic markers in our antibody panel, we manually gated 38 canonical immune cell populations (Figure S1B) and evaluated immune cell population frequencies, protein expression patterns, and immune cell signaling pathways specific to COVID-19 course escalation and resolution.

**Figure 1:**
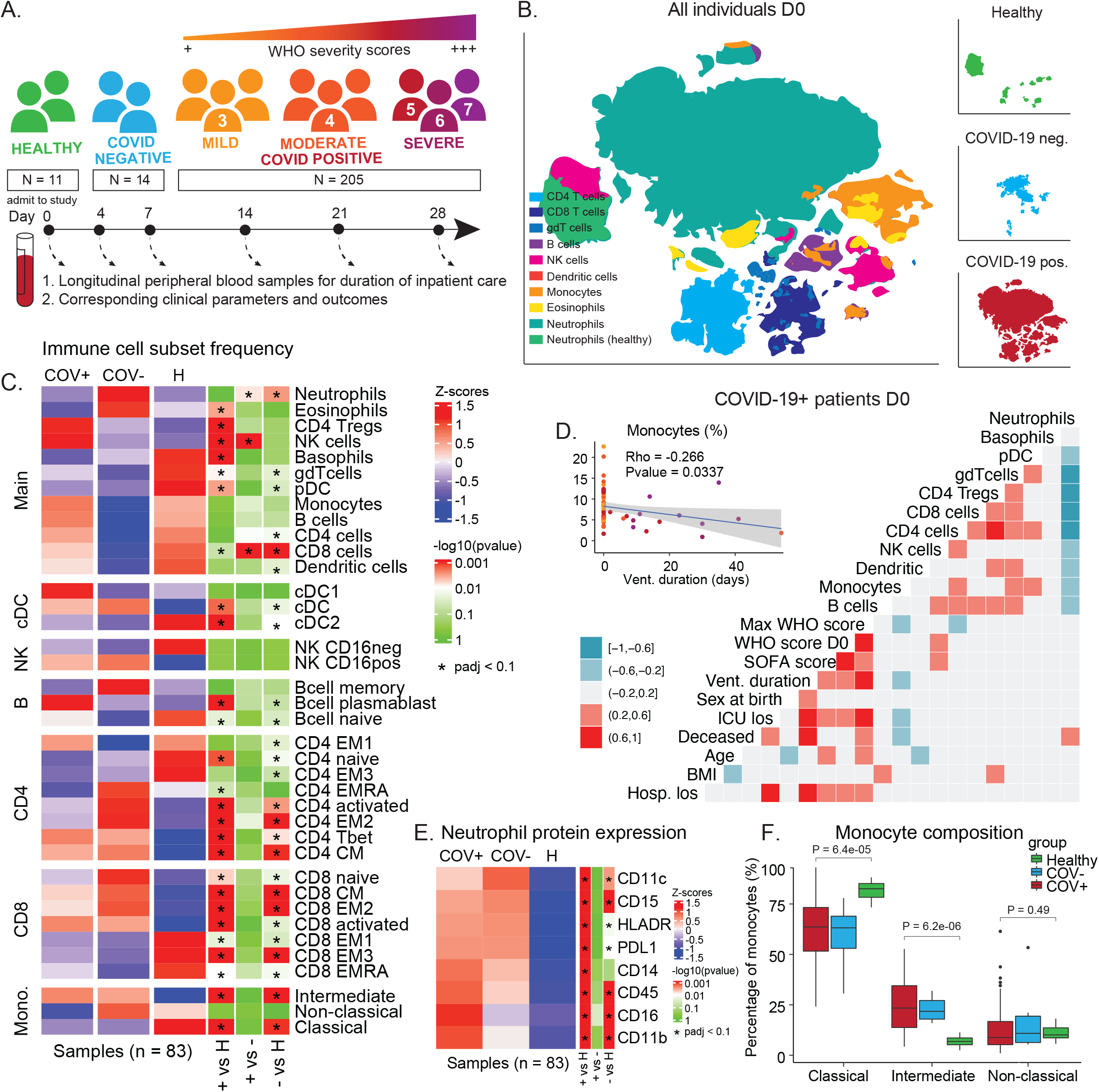
COVID-19 immune phenotype and composition is highly divergent from healthy individuals and has unique features compared to other severe respiratory infections. **A)** Overview of cohort. Patients were admitted to the hospital and enrolled in the study at D0. Peripheral blood samples were collected throughout the duration of stay. Corresponding clinical parameters and WHO scores were documented. 205 samples from 81 COVID-19 positive patients were included in the final cohort. Additionally, 14 samples from 7 COVID-19 negative patients with other respiratory diseases and 11 healthy individuals were included in the study. **B)** t-SNE plot of all patient samples at D0 (n = 83) using phenotypic markers colored by major immune cell populations. Upper right panel: t-SNE plot of healthy samples (n = 11); middle right panel: t-SNE plot of COVID-19 negative samples (n = 6); lower right panel: t-SNE plot of COVID-19 positive samples (n = 66). **C)** Immune cell population abundance at D0 in COVID-19 positive (+), COVID-19 negative (-) patients, and healthy individuals (H). P-values obtained by Wilcoxon Rank Sum Test, followed by Benjamini-Hochberg correction with FDR < 0.1. **D)** Correlation between cell population abundance at D0 and clinical outcomes, e.g. ventilation duration (vent_duration) and hospital length of stay (hosp_los) for COVID-19+ patients. Correlation estimates are obtained by Pearson correlation. **E)** Protein expression on neutrophils (F) in COVID-19 positive (COV+), COVID-19 negative (COV-) patients, and healthy controls at D0 (Wilcoxon Rank Sum Test, Benjamini-Hochberg correction with FDR < 0.1). **F)** Frequency of monocyte subsets in COVID-19 positive (COV+), COVID-19 negative (COV-) patients, and healthy controls at D0. P-values obtained by Wilcoxon Rank Sum Test.

### Unique immune cell compositions between COVID-19 patients, COVID-19 negative patients, and healthy individuals on day of admission

First, we characterized the immunological landscape of COVID-19 patients, COVID-19 negative patients (critically ill, mechanically ventilated controls), and healthy individuals to assess immunological signatures that may be unique to COVID-19 at day of admission (day 0; D0). Dimensionality reduction by t-SNE using only phenotypic markers revealed distinct immune cell compositions between COVID-19 positive, COVID-19 negative, and healthy individuals (Figure 1B). Consistent with previous studies, COVID-19 patients exhibited a significantly different immune cell composition compared to healthy individuals, with significant frequency changes across almost all manually gated immune cell populations (Figure 1C) (Mathew et al. 2020). To determine modules of immune changes, we evaluated if distinct immune cell populations correlate with each other as well as with patient demographics or clinical parameters. We found a coordinated adaptive immune response in which several T cell subsets and B cell frequencies were positively correlated with one another (Figure 1D). In contrast, the innate arm demonstrated a dichotomous relationship, with neutrophil and monocyte frequencies being anti-correlated. Additionally, monocyte frequencies at day 0 were positively correlated with T cell subsets and negatively correlated with ventilation duration (Figure 1D), suggesting there may be a coordinated immune response associated with better clinical outcome.

### Monocyte and neutrophil composition reveal unique compartmental shifts in innate immune arm of COVID-19 infection

Large shifts in innate immune compartments were evident between COVID-19 patients, patients with other respiratory infections, and healthy controls (Figure 1B); therefore, we further investigated the composition of neutrophils and monocytes. While neutrophil frequency was not significantly different between COVID-19 patients and the healthy individuals (Figure 1C and S1C), we found that a variety of proteins were altered in their expression on neutrophils across groups. Neutrophils from COVID-19 patients exhibited significantly increased expression of CD11c, CD14, CD16, and PD-L1, suggesting a highly activated and inflammatory neutrophil phenotype in COVID-19 patients (Figure 1E). Additionally, while the frequency of all monocytes was comparable between groups (Figure 1C), composition of monocyte subsets (defined as classical, intermediate, and non-classical) was significantly different between patients with COVID-19 and other respiratory infections compared to healthy individuals. Patients exhibited a significant increase in the frequency of intermediate monocytes along with a relative decrease in classical monocytes (Figure 1F).

### Cross-sectional analysis of COVID-19 severity groups reveals few immunological features that distinguish severity states, requiring a new approach to evaluating immune trajectories in our patient cohort

Having established the major differences between COVID-19 patients, COVID-19 negative patients, and healthy individuals at D0, we turned to evaluate the immunological differences between COVID-19 severity groups across time (Figure S1D). Surprisingly, we found no significant differences between severity groups at D0 and only few population differences at D4 and D7 (Figure S1E and S1F). Within each severity group, comparisons across time showed that plasmablasts contract from D0 to D7 in the majority of severe COVID-19 patients (Figure S1G), while CD4 activated T cells are upregulated from D0 to D7 in mild COVID-19 patients (Figure S1H). The paucity of differences between severity groups suggested that perhaps significant variability exists in the timing of disease escalation and resolution across individuals and therefore the immunological processes that mediate these changes over time.

### Early, coordinated, and activated immune cell signaling states in COVID-19 patients

To gain insights into key immune cell signaling modules associated with COVID-19, we measured the phosphorylation state of 14 signaling molecules across all immune cell subsets (Figure 2A). First, we evaluated the median level of phosphorylated signaling proteins across all CD45+ hematopoietic PB cells in COVID-19 positive, COVID-19 negative, and healthy individuals at D0. Differential expression analysis revealed five signaling molecules (pSTAT1, pPLCg2, pZAP70/pSyk, pCREB, and pSTAT3) that were upregulated in COVID-19 patients compared to healthy individuals (Figure 2B). To determine if a specific cell type was driving the higher signaling state in COVID-19 patients, we evaluated the median phosphorylation state of the respective signaling molecules within all manually gated immune cell subsets. We found significantly higher median signaling across the majority of cell subsets, showing that immune cell signaling states are coordinated across most cell types simultaneously and not driven by signaling within a specific cell type (Figure S2A).

**Figure 2:**
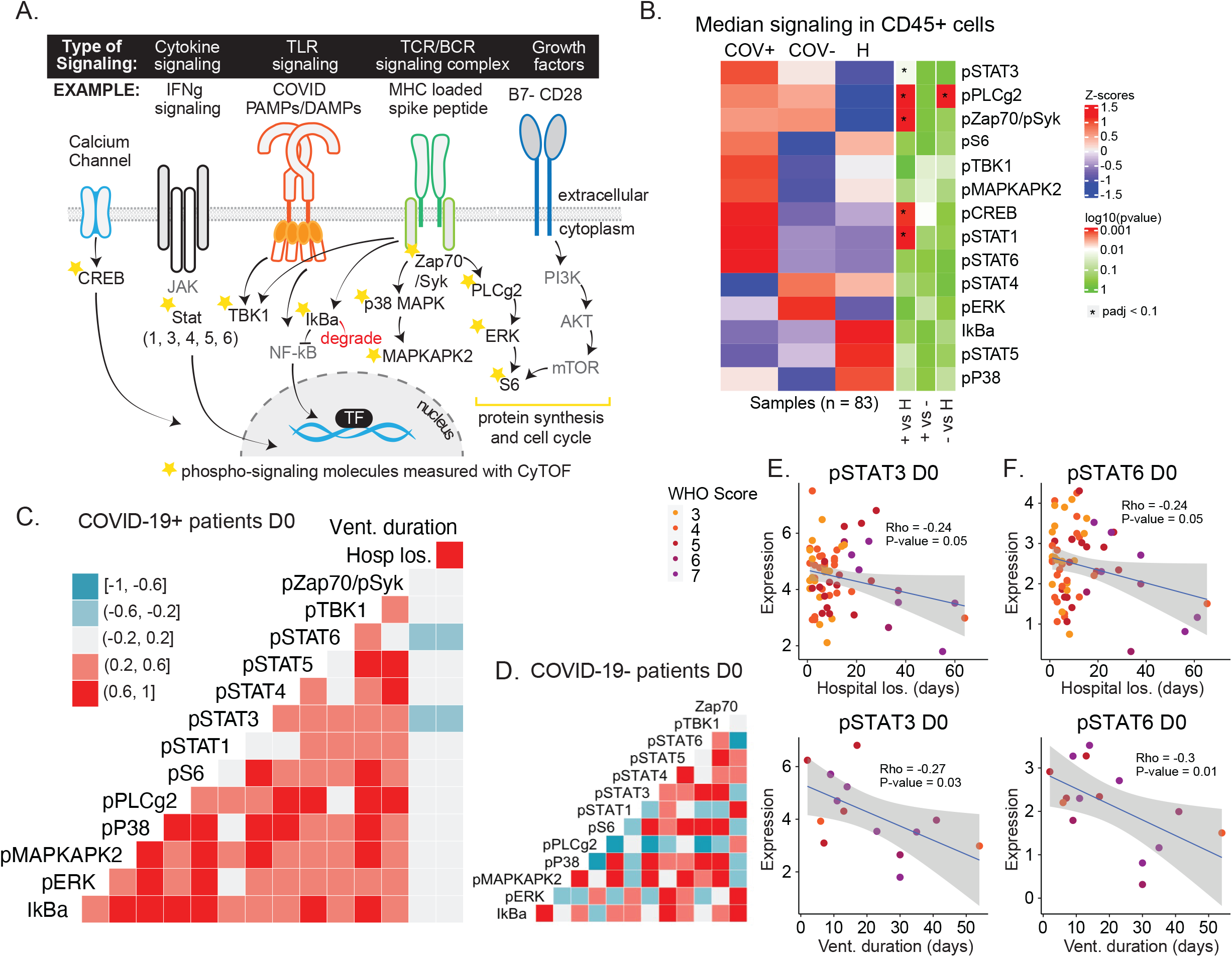
Early coordinated immune signaling defines COVID-19 patients with high pSTAT3 and pSTAT6 associated with favorable clinical outcomes. **A)** Signaling schematic. Stars denote phosphorylated signaling molecules that are measured in the CyTOF panel. **B)** Expression of signaling molecules in CD45+ CD235a/b-negative peripheral blood immune cells at D0 in COVID-19 positive (+), COVID-19 negative (-) patients, and healthy individuals (H). P-values obtained by Wilcoxon Rank Sum Test, followed by Benjamini-Hochberg correction with FDR < 0.1. **C+D)** Correlation between signaling molecule expressions at D0 and clinical outcomes, e.g. ventilation duration (vent_duration) and hospital length of stay (hosp_los) for COVID-19+ patients (C) and COVID-19-patients (D). Correlation estimates are obtained by Pearson correlation. **E+F)** Correlation between pSTAT3 (E) or pSTAT6 (F) signaling at D0 and hospital length of stay or ventilation duration. Correlation estimates and p-values are obtained by Pearson correlation.

To investigate coordinated signaling modules in CD45+ cells, we correlated the expression of signaling molecules at D0. For COVID-19 patients, we observed a coordinated positive signaling response (Figure 2C), while this coordination was absent in patients with other respiratory infections or sepsis (Figure 2D). Additionally, we correlated signaling molecule expression to hospital length of stay and ventilation duration (Figure 2C) and found significant negative correlations between these parameters and levels of pSTAT3 (Figure 2E) and pSTAT6 (Figure 2F), suggesting that higher pSTAT3 and pSTAT6 signaling at time of admission corresponds to better clinical outcomes. Finally, we evaluated signaling differences within and across severity groups at D0, D4, and D7, but observed no significant changes (Figure S2B and S2C).

### Conserved immunological processes and changes in cell signaling states accompany disease resolution and discharge

Although cross-sectional analysis can provide insights into the immunological state of COVID-19 patients and severity groups, the natural heterogeneity of patient immune responses and significant differences in their disease time courses may obscure immunological processes that mediate recovery. Therefore, we aimed to identify conserved changes within patients, over time, that are tied to clinically relevant outcomes. Given that the majority of our patients successfully recovered from the infection, albeit after differing lengths of hospitalization, we investigated immunological changes that occurred within patients from time of admission (tp1) to time of discharge (tp2) from the hospital (n = 32) (Figure 3A and S3A). For this analysis, we included patients who were discharged within 30 days of admission across all disease severity states at time of enrollment, allowing us to identify conserved features among all COVID-19 patients who successfully recover. A variety of immune cell subsets significantly changed in frequency between tp1 and tp2 (Figure 3B). Monocytes as well as activated CD4 and CD8 T cells significantly increased at the time of discharge (tp2) as patients resolved the infection (Figure 3C). Conversely, neutrophils and conventional type 1 dendritic cells (cDC1s) significantly decreased in frequency by time of discharge (Figure 3C). For most COVID-19 patients, the overall composition of immune cells became more similar to that of healthy individuals at the time of discharge compared to the time of enrollment (Figure 3D). However, some immune cell populations exhibited deviations away from healthy at the time of discharge, most notably activated CD4 and CD8 T cells (CD38+ HLA-DR+) as well as monocytes (Figure 3E). This indicates that the immune state at the time of discharge is characterized by the restoration of certain elements of the immune response that were perturbed early in infection alongside a continued immunological process that proceeds past the time patients stabilize for discharge.

**Figure 3:**
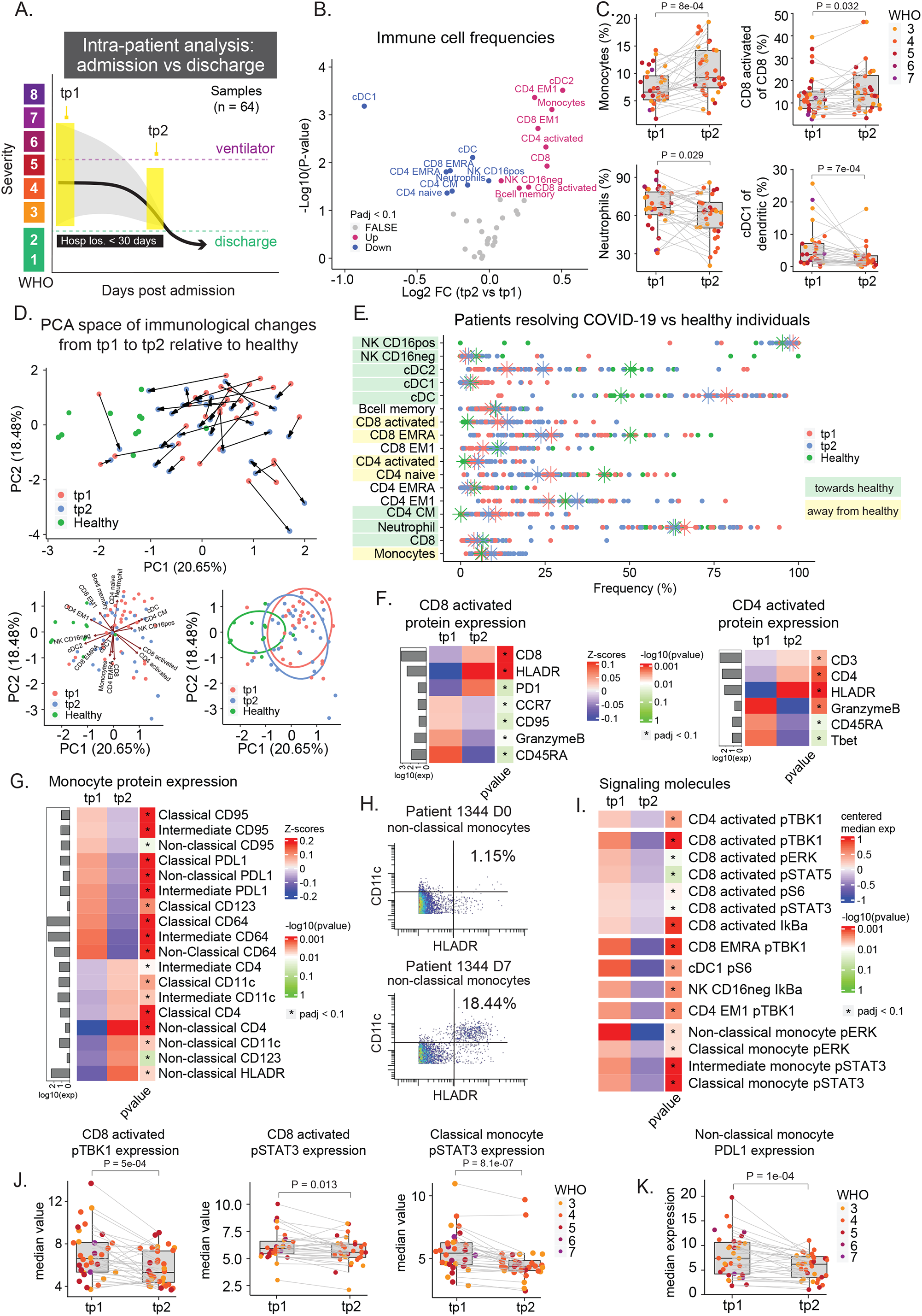
Conserved immunological processes accompany COVID-19 resolution and hospital discharge. **A)** Illustration of intra-patient analysis from admission to discharge for patients who are successfully discharged from the hospital within 30 days of admission (n = 32). **B)** Paired differential expression analysis of immune cell populations between the first (tp1) and second (tp2) timepoints illustrated in 3A (paired Wilcoxon Rank Sum Test). The log2 fold changes (tp2 vs tp1) are plotted against the negative log10(p-values). Colors indicate if cell populations are significantly down-(blue) or upregulated (purple) from tp1 to tp2 or not differentially expressed (FALSE, grey) after Benjamini-Hochberg correction, FDR < 0.1. **C)** Frequency of monocytes, neutrophils, cDC1, and CD8 activated T cells at tp1 and tp2. Lines connect samples from the same patient. P-values obtained by paired Wilcoxon Rank Sum Test. CD8 activated T cells and cDC1 cells are shown as a percentage of parent populations (e.g. CD8 T cells and dendritic cells, respectively), while monocytes and neutrophils are shown as a percentage of all cells. **D)** Principal component analysis of significant immune cell subsets in 3B for tp1, tp2, and healthy controls. Immune cell directionality and contribution to PCA space denoted on right (top). Summary ellipsoid of tp1, tp2, and healthy patients in PCA space on right (bottom). **E)** Population frequencies of significant immune cell subsets in 3B for tp1, tp2, and healthy controls. Stars indicate median value for each group. Cell populations are highlighted in green if tp2 is closer to healthy than tp1, and highlighted in yellow if tp2 is moving away from healthy. **F+G)** Protein expression on CD8- and CD4 activated T cells (F) and on monocyte subsets (G) at tp1 and tp2. Mean protein expression values have been log10 transformed, scaled, and centered on heatmap. Bars indicate mean protein expression across all samples. Only significant proteins are shown (Wilcoxon Rank Sum Test, Benjamini-Hochberg correction with FDR < 0.1). **H)** Scatter plots of CD11c and HLA-DR expression on non-classical monocytes in patient 1344 at D0 (top) and D7 (bottom). **I)** Expression of signaling molecules in significant immune cell subsets in 3B at tp1 and tp2. Median signaling expression values have been centered on heatmap. Only significant signaling molecules are shown (Wilcoxon Rank Sum Test, Benjamini-Hochberg correction with FDR < 0.1). **J)** Expression of pTBK1 in CD8 activated T cells, and pSTAT3 expression in CD8 activated T cells and classical monocytes at tp1 and tp2. Lines connect samples from the same patient. P-values obtained by paired Wilcoxon Rank Sum Test. **K)** Expression of PDL1 on non-classical monocytes at tp1 and tp2. Lines connect samples from the same patient. P-values obtained by paired Wilcoxon Rank Sum Test.

### Patients who successfully resolve COVID-19 have robust pan-hematopoietic signaling and cytotoxic activated T cells at day 0

To obtain more granular insights into the immunological perturbations that accompany COVID-19 recovery, we evaluated phenotypic changes and signaling dynamics within immune cell populations that changed during disease resolution. We focused on cell populations whose frequencies move away from levels observed in healthy controls, indicating they continue to have a dynamic response during infection resolution. Activated CD4 and CD8 T cells exhibited a reduction in the expression of GranzymeB and CD45RA as patients transition from early infection to discharge (Figure 3F and S3B), consistent with a transition from more activated effector cells to more of a memory phenotype. Interestingly, we also observed a significant change in the phenotype of circulating monocytes, which expressed high levels of PD-L1 at time of admission but higher levels of CD4, CD11c, and HLA-DR at time of discharge (Figure 3G, 3H, and S3B). Similarly, we observed a reduction in PD-L1 expression on neutrophils at time of discharge (Figure S3B).

We then analyzed the median values of phosphorylated signaling molecules within the relevant immune cell subtypes to evaluate changes in cell signaling during this resolution phase. A variety of cell signaling proteins were significantly downregulated within the key immune cell populations at time of discharge (Figure 3I). Several signaling molecules changed in a coordinated fashion across different immune cell types (e.g. pTBK1, pERK, and pSTAT3), with the broadest signaling changes observed in activated CD8 T cells and monocyte subsets (Figure 3I and 3J). These observations are consistent with previous studies describing the relationship between IL-6 expression and pSTAT3 signaling and subsequent upregulation of PD-L1 in monocytes (Figure 3F and 3K) (W. Zhang et al. 2020). Although signaling trajectories trended in the same direction among most patients (Figure S3C), we did not observe a clear trend towards healthy individuals (Figure S3D), likely explained by the expression variability and difficulty of measuring signaling molecules in rare populations of healthy cohorts e.g. activated CD8 T cells (Figure 3E). Taken together, our results suggest that a coordinated set of changes in immune cell abundances and signaling states occur in patients who successfully resolve COVID-19.

### Immune features associated with COVID-19 resolution are absent in patients who are hospitalized for more than 30 days or die from COVID-19

To determine if the immune features identified in the resolution phase are specific to patient recovery, we analyzed patients who had delayed disease resolution, i.e. who remained hospitalized for more than 30 days (“late discharge”; n = 6 patients) or who died from COVID-19 (“ultimately deceased”; n = 5 patients) (Figure 4A and S4A). First, we evaluated the immune cell population changes occurring within these patients over a similar period from the time of admission, but found no significant changing populations for either group (Figure S4B). We asked if the lack of immune remodeling between these timepoints was due to a reflection of an insufficient initial response or, alternatively, a sustained immune response that failed to resolve. In fact, both baseline immune cell frequencies at the time of admission and the magnitude of their changes were different, though in different ways for different elements of the immune response (Figure 4B, S4C, S4D, and S4E). In deceased patients, neutrophil frequencies were excessively elevated at both tp1 and tp2, while monocytes started at a lower frequency and failed to reach levels comparable to resolving patients (Figure 4B and S4F). Activated CD8 T cells were present at similar abundances across groups at the time of admission but became much more abundant in late discharge and ultimately deceased patients (Figure 4B and 4C). In contrast, activated CD4 T cells were already more elevated in late discharge and ultimately deceased patients at the time of admission and became even more elevated over time (Figure 4B). An increase in the abundance of cDC1s was notably absent in ultimately deceased patients at the time of admission, while they were substantially more elevated in late discharge patients (Figure 4B).

**Figure 4:**
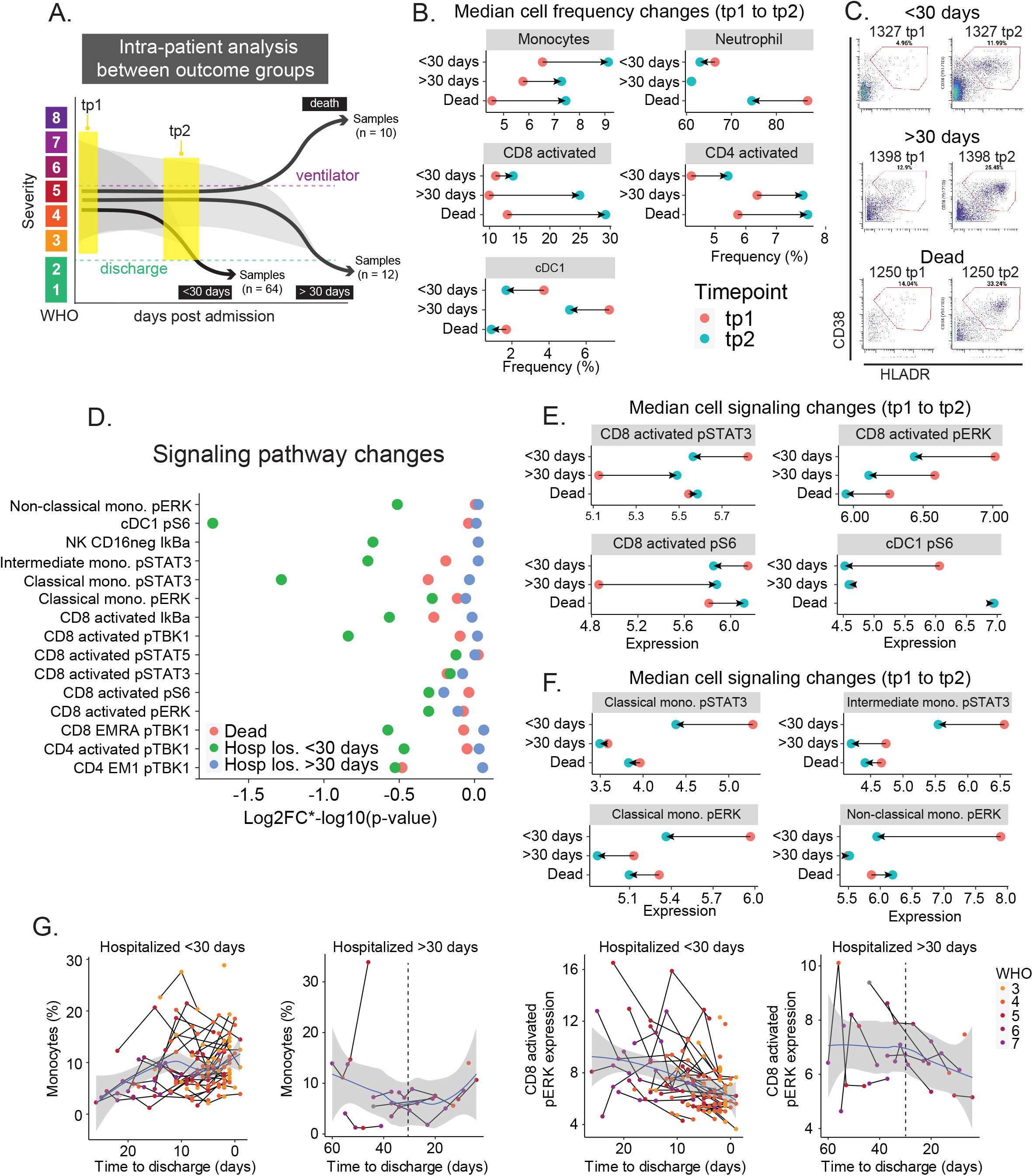
Immune features associated with COVID-19 resolution are absent in patients who are discharged late or die from COVID-19. **A)** Illustration of intra-patient analysis of patients who are hospitalized for >30 days (n = 6) and patients who die (n = 5). **B)** Median cell population frequencies at tp1 (red) and tp2 (blue) for patients who are discharged <30 days, >30 days, and deceased. **C)** Representative scatter plots of activated CD8 T cells (defined by CD38 and HLA-DR expression), at tp1 (left) and tp2 (right) for patients who are discharged <30 days, >30 days, and deceased. **D)** Magnitude of change illustrated by log2FC*-log10(pvalue) of signaling molecules (identified in Figure 3I) for patients who are discharged within 30 days (<30 days, green), discharged after 30 days (>30 days, blue), and die (red). P-values obtained by paired Wilcoxon Rank Sum Test. **E+F)** Median signaling molecule expression at tp1 (red) and tp2 (blue) for patients who are discharged <30 days, >30 days, and deceased. **G)** Monocyte frequencies (left plots) and CD8 activated pERK expressions (right plots) relative to time to discharge in all samples from patients who are discharged <30 days (n = 142 samples) or >30 days (n = 30 samples). Black lines connect samples from the same patient. Blue lines and grey shadows represent the best fitted smooth line and 95% confidence interval. Dotted lines intersect the x-axis at day 30.

### Elevated cell signaling at time of admission is associated with COVID-19 resolution

Next, we evaluated signaling dynamics in late discharge and ultimately deceased patients to determine if observed changes in cell frequencies were accompanied by dysfunctional signaling. In contrast to patients resolving COVID-19 in <30 days, which exhibited consistent changes from high to low signaling states over time, we observed no significant changes for late discharged and ultimately deceased patients (Figure 4D, S4G, S4H, and S4I). Instead, these patients exhibited discoordinate signaling directionality in activated CD8 T cells (Figure 4E), a complete lack of pS6 signaling in cDC1 cells (Figure 4E), and less signaling at tp1 across monocyte subsets (Figure 4F). Interestingly, when the late discharged patients are within 30 days of discharge, the trajectory of several immune resolution features, e.g. monocytes, neutrophils, and signaling molecules, resembles the recovery trajectories in patients hospitalized <30 days, suggesting that the resolution phase engages in these patients as well before they are discharged (Figure 4G and S4K). Taken together, these results indicate that late discharge and ultimately deceased patients exhibit reduced immune cell signaling at the time of hospitalization. While some of these cell signaling pathways became elevated at later time points in these patients, others were not changing at all. Furthermore, these results suggest that the immune processes observed during resolution through discharge are specific to a successful response against COVID-19.

### Core immune resolution features characterize COVID-19 patients recovering from ventilation

Having established immune features that accompany COVID-19 resolution among our entire patient cohort, we next examined the immunological changes within only the most severe patients who required mechanical ventilation (Figure S5A). We analyzed immunological changes between three key time points; the first time point after a patient was intubated (tp1), the last time point before they were extubated (tp2), and the first time point after a patient was successfully extubated (tp3) (Figure 5A). This allowed us to evaluate the immunological dynamics that occur during ventilation (tp1 vs tp2), and during successful recovery from intubation (tp1 vs tp3). First, we analyzed the within-patient immune cell frequency changes between tp1 and tp3 (n = 9, S5B). Consistent with patients resolving COVID-19, monocytes and activated CD4 and CD8 T cells significantly increased in frequency, while neutrophil frequency decreased during ventilation resolution (Figure 5B and 5C). Additionally, ventilation resolution was characterized by an increase of CD4 regulatory T cells (Tregs) and basophils at time of recovery (Figure 5B). These changes collectively were associated with a coordinated trajectory of recovery from tp1 to tp3 (Figure 5D). Despite these coordinated changes, patients did not return to an immune composition comparable to healthy donors, indicating that the time of extubation remains an active immunological phase of disease resolution from the most severe form of COVID-19. Some key immune cell populations that remain different from healthy controls included both activated CD4 and CD8 T cells as well as Tregs (Figure 5E and S5C). Of these changes, only the observed increase in activated CD8 T cells was apparent within patients during intubation (tp1 vs tp2), suggesting that additional dynamic changes are specific to the resolution of severe COVID-19 (Figure S5D and S5E).

**Figure 5:**
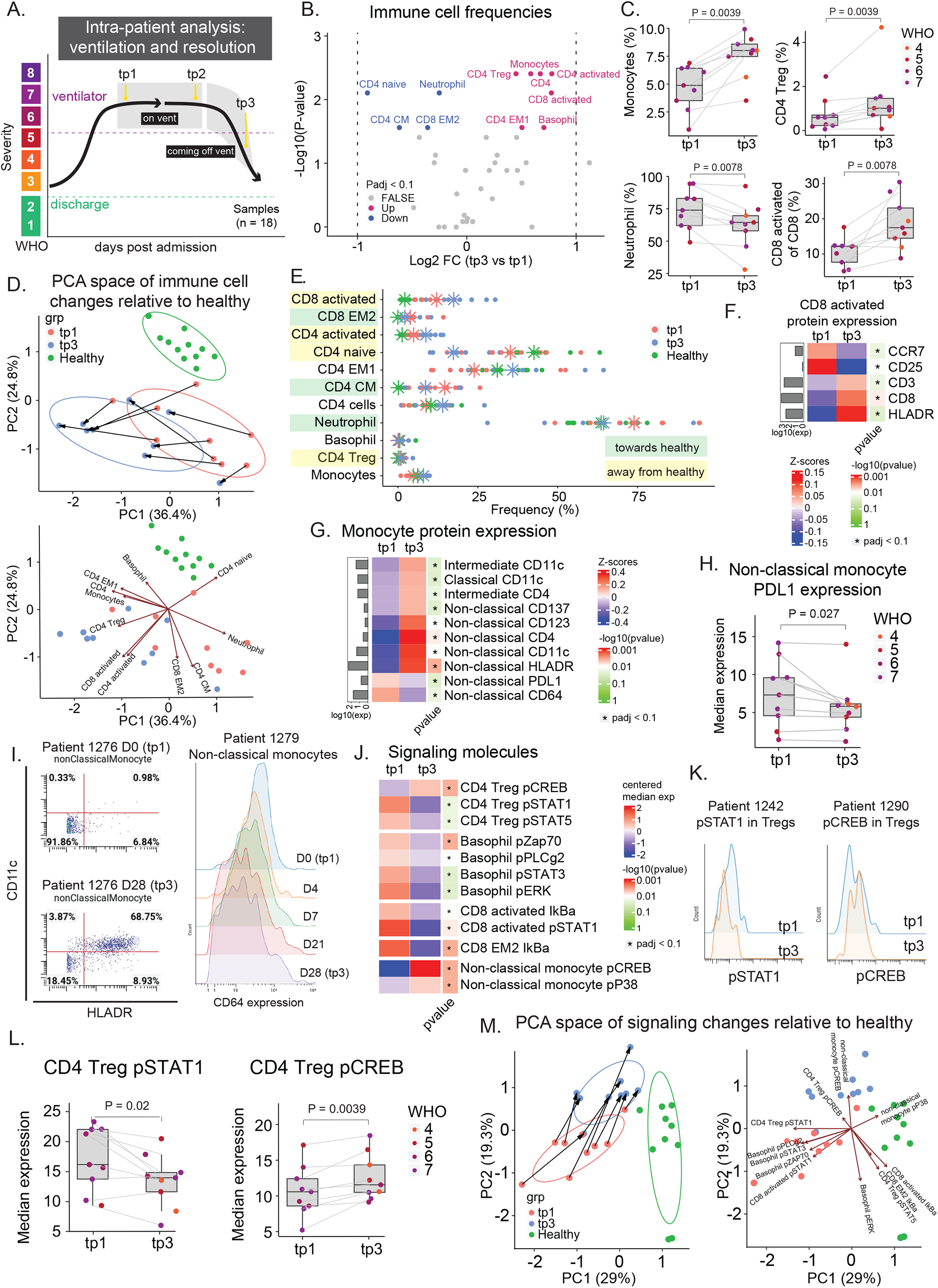
Recovery from severe COVID-19 requires core immune resolution features and additional regulatory T cell and basophil upregulation. **A)** Illustration of intra-patient analysis of ventilated patients. Three timepoints are considered: tp1 (first sample after a patient has been put on a ventilator), tp2 (last sample before the patient is removed from a ventilator), and tp3 (first sample after a patient is successfully removed from ventilation support). **B)** Paired differential expression analysis of immune cell populations between the first (tp1) and third (tp3) timepoints illustrated in 5A (paired Wilcoxon Rank Sum Test). The log2 fold changes (tp3 vs tp1) are plotted against the negative log10(p-values). Colors indicate if cell populations are significantly down- (blue) or upregulated (purple) from tp1 to tp3 or not differentially expressed (FALSE, grey) after Benjamini-Hochberg correction, FDR < 0.1. **C)** Frequency of monocytes, neutrophils, CD4 Treg, and CD8 activated T cells at tp1 and tp3. Lines connect samples from the same patient. P-values obtained by paired Wilcoxon Rank Sum Test. CD8 activated T cells are shown as a percentage of parent population (e.g. CD8 T cells), while monocytes, neutrophils, and CD4 Tregs are shown as a percentage of all cells. **D)** Principal component analysis of significant immune cell subsets in 5B for tp1, tp3, and healthy controls. Immune cell directionality and contribution to PCA space denoted on the right. **E)** Population frequencies of significant immune cell subsets in 3B for tp1, tp3, and healthy controls. Stars indicate median value for each group. Cell populations are highlighted in green if tp3 is closer to healthy than tp1, and highlighted in yellow if tp3 is moving away from healthy. **F+G)** Protein expression on CD8 activated T cells (F) and on monocyte subsets (G) at tp1 and tp3. Mean protein expression values have been log10 transformed, scaled, and centered on heatmap. Bars indicate mean protein expression across all samples. Only significant proteins are shown (Wilcoxon Rank Sum Test, Benjamini-Hochberg correction with FDR < 0.1). **H)** Expression of PDL1 on non-classical monocytes at tp1 and tp3. Lines connect samples from the same patient. P-values obtained by paired Wilcoxon Rank Sum Test. **I)** Left: Scatter plots of CD11c and HLA-DR expression on non-classical monocytes in patient 1276 at D0 (tp1, top) and D28 (tp3, bottom). Right: Expression of CD64 on non-classical monocytes for patient 1279 from D0 (tp1) to D28 (tp3). **J)** Expression of signaling molecules in significant immune cell subsets in 5B at tp1 and tp3. Median signaling expression values have been centered on heatmap. Only significant signaling molecules are shown (Wilcoxon Rank Sum Test, Benjamini-Hochberg correction with FDR < 0.1). **K)** Expression of pSTAT1 (left) and pCREB (right) in CD4 Tregs at tp1 (blue) and tp3 (orange) for representative patients. **L)** Expression of pSTAT1 and pCREB in CD4 Tregs at tp1 and tp3. Lines connect samples from the same patient. P-values obtained by paired Wilcoxon Rank Sum Test. **M)** Principal component analysis of significant signaling molecules in 5I for tp1, tp3, and healthy controls. Immune cell directionality and contribution to PCA space denoted on the right.

### COVID-19 ventilation recovery is associated with T cell and monocyte phenotypic changes and a transition from pSTAT to pCREB dominated signaling

Next, we further analyzed changes in immune cell activation and cell signaling dynamics that accompany ventilation resolution. Consistent with recovery trajectories in patients resolving COVID-19, activated CD8 T cells expressed higher levels of HLA-DR and lower levels of CCR7 at the time of extubation (Figure 5F), while neutrophils expressed lower levels of PD-L1 (Figure S5F). Additionally, while there was no difference in monocyte subset frequencies (Figure S5G), non-classical (CD16+) monocytes exhibited a shift from a CD64+ PD-L1+ phenotype during ventilation to a CD4+ CD11c+ HLA-DR+ activated monocyte phenotype at the time of extubation (Figure 5G, 5H, and 5I). CD64+ expression on non-classical monocytes were incrementally decreased between tp1 and tp3 demonstrating a progressive downregulation during the resolution phase (Figure 5I).

Cell signaling states also changed markedly from the time of intubation to the time of extubation. During early time points of mechanical ventilation (tp1), higher levels of pSTAT1, pSTAT3, and pSTAT5 signaling was evident in CD4 Tregs, basophils, and activated CD8 T cells (Figure 5J, 5K, 5L, and S5H). Conversely, pCREB signaling was significantly increased after extubation (tp3) in CD4 Tregs and non-classical monocytes (Figure 5J, 5K, 5L, and S5H), suggesting there is a transition from inflammatory cytokine signaling response to pro-survival signaling within these cells, specifically. Visualizing these signaling trajectories in PCA space revealed a coordinated trajectory of immune cell signaling that accompanies extubation across patients (Figure 5M), though signaling states remained distinct from those in healthy individuals (Figure S5I). Taken together, our analyses identify a conserved set of immunological processes that are consistent among patients who recovered from mechanical ventilation as a result of COVID-19, elucidating an additional layer of immunological changes that are unique to these patients compared to recovery in patients who did not require mechanical ventilation.

### Core immune resolution features define patients with better clinical outcomes at time of admission

Having identified a signature of immune remodeling during COVID-19 recovery, we next investigated if the early presence of these features were associated with better patient outcomes. We evaluated the immune composition of severe COVID-19 patients before or on the day they were ventilated (vent, n = 13) and compared it to the immunological state at time of admission (D0) for patients who never required ventilation (no vent, n = 50) (Figure 6A and S6A). Differential abundance analysis of immune cell frequencies revealed higher frequencies of monocytes and CD4 Tregs, as well as decreased neutrophil frequencies, in patients who never required ventilation (Figure 6B and 6C). Similar results were obtained when exclusively analyzing samples collected prior to ventilation (vent, n = 8) (Figure S6B and S6C). Patients who never required ventilation exhibited an immune state more similar to those of the healthy controls (Figure S6D). While monocytes were significantly downregulated at time of admission in patients who required ventilation, we observed a consistent increase from time of intubation to time of discharge with the highest incline occurring right after time of extubation (Figure 6D). The opposite directionality was observed for neutrophils (Figure 6D). Interestingly, CD4 Tregs, which are known to play a role in ARDS resolution and pulmonary recovery, demonstrate a gradual increase in frequency during patient intubation followed by the steepest increase after extubation (Mock et al. 2014; Garibaldi et al. 2013) (Figure S6E). Additionally, the phenotype of monocytes in patients who never require ventilation resembles the activated monocyte subset identified during discharge and ventilation recovery, expressing significantly higher levels of CD4 and CD11c (Figure S6F and S6G). Furthermore, basophil and CD4 Treg signaling states that were identified during ventilation resolution were already significantly higher in patients who required ventilation at time of admission (Figure 6E and S6H) and consistently decreased during ventilation (Figure 6F). Taken together, our results show a set of conserved core immune features that accompany disease resolution with additional features that identify patients who recover from ventilation (Figure 6G). These ventilation specific features are significantly different at time of admission between patients who will require mechanical ventilation and those that never require ventilation, and thus associated with poorer clinical outcomes (Figure S6I).

**Figure 6:**
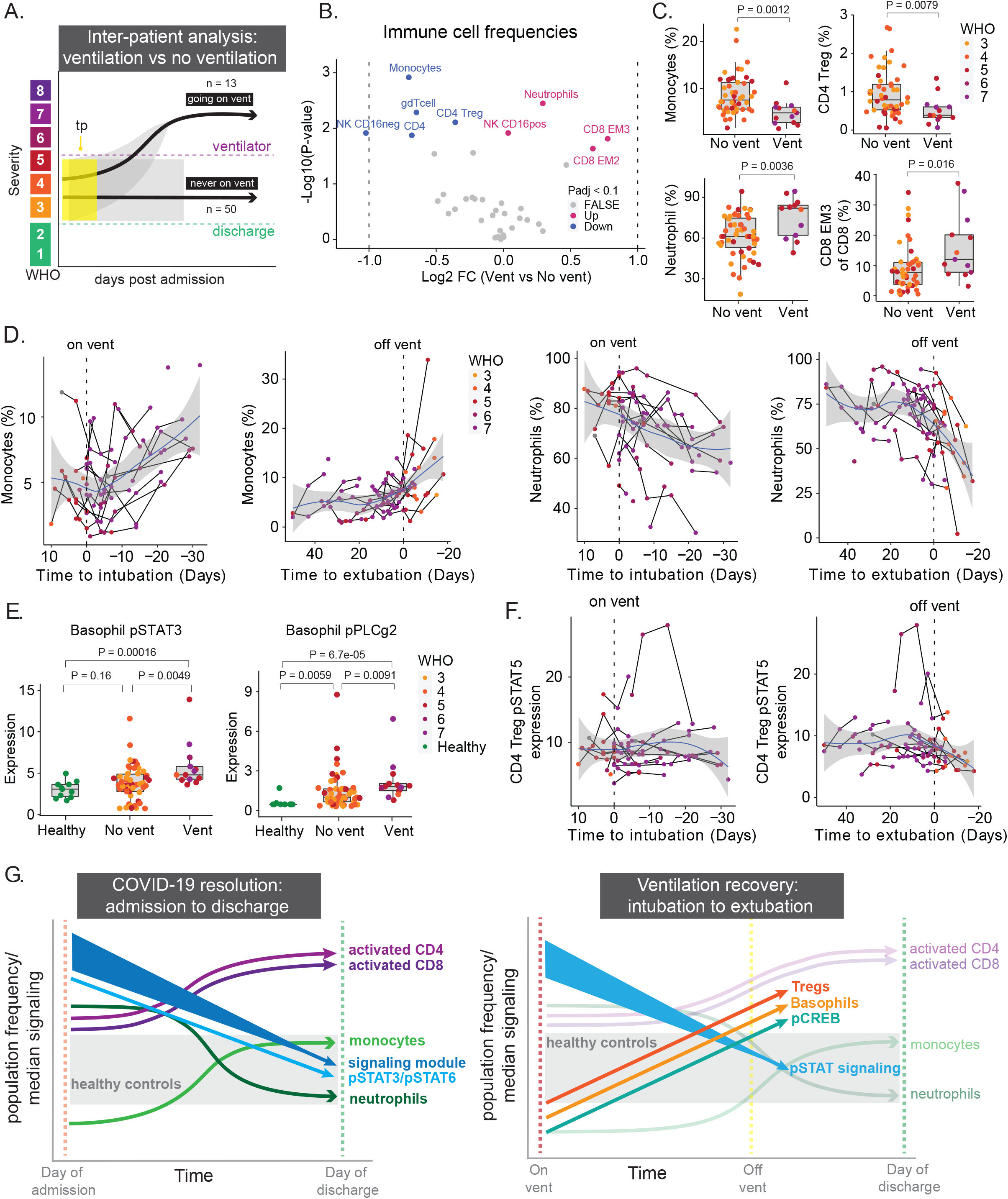
Core immune resolution features define patients with better clinical outcomes at time of admission. **A)** Illustration of inter-patient analysis of ventilated patients (vent, n = 13) vs patients who are never ventilated (no vent, n = 50). For ventilated patients, the latest sample before the patient is put on a ventilator is used. For non-ventilated patients, D0 is used. **B)** Differential expression analysis of immune cell populations between ventilated and non-ventilated patients illustrated in 6A (Wilcoxon Rank Sum Test). The log2 fold changes (vent vs no vent) are plotted against the negative log10(p-values). Colors indicate if cell populations are significantly down- (blue) or upregulated (purple) for vent vs no vent or not differentially expressed (FALSE, grey) after Benjamini-Hochberg correction, FDR < 0.1. **C)** Frequency of monocytes, neutrophils, CD4 Tregs, and CD8 EM3 T cells in vent and no vent patients. P-values obtained by Wilcoxon Rank Sum Test. CD8 EM3 T cells parent population (e.g. CD8 T cells), while monocytes, neutrophils, and CD4 Tregs are shown as a percentage of all cells. **D)** Monocyte (left plots) and neutrophil (right plots) frequencies relative to intubation / extubation in all samples from ventilated patients. Black lines connect samples from the same patient. Blue lines and grey shadows represent the best fitted smooth line and 95% confidence interval. Dotted lines intersect the x-axis at day of intubation / extubation. **E)** Expression of pSTAT3 and pPLCg2 in basophils in non-ventilated and ventilated patients as well as healthy individuals. P-values obtained by Wilcoxon Rank Sum Test. **F)** Expression of pSTAT5 in CD4 Tregs relative to intubation / extubation in all samples from ventilated patients. **G)** Graphical summary depicting the trajectories of key immune features involved in COVID-19 resolution and ventilation recovery.

## Discussion

Human immunology studies are inherently challenging because of the variability in baseline immune cell compositions, heterogeneity in immune responses, and difficulty in collecting longitudinal samples to track individuals over time. Because of the urgency to understand and respond to COVID-19, this cohort of patients provided a unique opportunity to recruit, study, and analyze a large number of individuals responding to the same infection over a finite period of time (April 2020 - April 2021). Since individuals recover from their infection across a variable amount of time, these studies highlight the benefit of longitudinal analysis anchored on key clinical events in the disease process. This analytical approach revealed the unifying trends among patients that define clinically relevant events such as discharge from the hospital or extubation after mechanical ventilation, regardless of initial disease severity or time to recovery.

Our findings are consistent with several recent reports of immune responses to COVID-19 while contributing a new understanding of the immunological processes that accompany disease recovery, including changes in immune cell signaling states. Although some studies have suggested that early intervention to modulate immune hyperactivation may be beneficial in severe COVID-19 (Lucas et al. 2020), these data indicate that early immune cell signaling, particularly pSTAT3 and pSTAT6, correlates with shorter hospitalization and ventilation duration. This indicates that an early robust immune response, driven by pSTAT signaling, and subsequent contraction during recovery may be beneficial to resolve COVID-19. In patients who require mechanical ventilation, additional immunological processes involving increased Tregs and basophils also accompany recovery in addition to the core recovery trajectory observed in patients who did not require ventilation. Additionally in our analysis, the STAT1 pathway downstream of type I IFN signaling was not differentially activated between patients with different disease severities. Instead, our study identified that many signaling pathways are activated simultaneously at the time of hospitalization, consistent with a recent report of concordant production of cytokines associated with type 1, 2, and 3 immune responses in patients with severe COVID-19 (Lucas et al. 2020). Despite the importance of B cells to generate SARS-CoV-2 neutralizing antibodies (Lucas et al. 2021), interestingly, our work did not identify changes in circulating B cells associated with the recovery trajectory. This finding aligns with the clinical observation that B cell deficient patients or patients with agammaglobulinemia can recover from COVID-19 (Bange et al. 2021; Soresina et al. 2020), and suggests that B cells may play a role in contributing to immunological memory as compared to the resolution of severe COVID19. Our work identified regulatory T cells as significantly changing only in patients who require ventilation, starting at significantly lower frequencies than in patients who never require ventilation support but gradually progressing to a steep increase after extubation. These findings are consistent with their critical role in pulmonary repair and ARDs recovery and specifically identify them as mediators in recovery from severe COVID-19 (Mock et al. 2014; Garibaldi et al. 2013).

Overall, our study provides an understanding of the core immunological changes that accompany disease recovery from severe COVID-19 and provides a foundational model of a successful anti-SARS-CoV-2 immune response. This working model of a recovering immune response trajectory provides a benchmark to contextualize divergent immune processes during poor disease outcomes in immunosuppressed or immunocompromised patients, long-haul COVID-19 patients, pediatric patients with MIS-C, or response to new variants. By elucidating a conserved trajectory of successful recovery, this study also nominates key immunological processes that could be targeted to enable recovery of severe disease in COVID-19 patients and perhaps other acute respiratory infections.

## Acknowledgments

This work was supported by generous grants from The Carlsberg Foundation (T.L.H.O.) and a COVID-19 Fast Grant (M.H.S.). C.E.B was supported by fellowships from the NCI (1F31CA260938-01), NSF Graduate Research Fellowship Program (GRFP), and the UCSF Discovery Fellowship. We would like to thank the NIAID Immunophenotyping Assessment in a COVID-19 Cohort (IMPACC) Network and the National Institutes of Health for their support (3U19AI077439-13S1 and 3U19AI077439-13S2). P.M.S was supported by the Howard Hughes Medical Institute through the James H. Gilliam Fellowships for Advanced Study program. C.S.C. was supported by NHLBI (R35 HL140026). C.R.L. has received funding from NHLBI and NIAID and C.M.H was supported by an NHLBI K23 and a DOD grant. Additionally, this project has been made possible in part by grant numbers 2019-202665 from the Chan Zuckerberg Foundation and TSK-020586 from Genentech. We acknowledge the Parnassus Flow Cytometry CoLab Facility supported in part by NIH Grants P30DK063720, S10OD018040, S10OD018040, and S10OD021822. M.H.S. is a Chan Zuckerberg Biohub investigator and a Parker Institute for Cancer Immunotherapy investigator.

## Author contributions

C.E.B, I.T, K.M.A, D.J.E, and M.H.S. conceived the project and designed CyTOF experiments. C.E.B, T.L.H.O, and M.H.S. conceptualized the study. C.E.B, I.T, D.M.M., and S.T. performed CyTOF experiments. T.L.H.O and C.E.B performed data analysis and generated all figures. C.E.B, T.L.H.O, and M.H.S. wrote the manuscript. M.H.S. supervised the study. C.S.C., C.M.H., C.R.L., M.F.K, P.G.W., D.J.E., founded and led the COMET Consortium. All authors read and approved the final manuscript.

## *The UCSF COMET Consortium

Ravi Patel^3,4^, Yumiko Abe-Jones^1^, Saurabh Asthana^2,3,4^, Alexander Beagle^5^, Sharvari Bhide^6^, Cathy Cai^7^, Maria Calvo^6^, Sidney A. Carrillo^8^, Suzanna Chak^8^, Zachary Collins^2,3,4^, Spyros Darmanis^9^, Gabriela K. Fragiadakis^3,4,10^, Rajani Ghale^8^, Jeremy Giberson^6^, Pat Glenn^11^, Ana Gonzalez^6^, Kamir Hiam-Galvez^12^, Alejandra Jauregui^8^, Serena Ke^6,13^, Tasha Lea^2^, Deanna Lee^6,13^, Raphael Lota^11^, Leonard Lupin-Jimenez^3^, Viet Nguyen^6,13^, Nishita Nigam,^1^ Logan Pierce^1^, Priya Prasad^1^, Arjun Rao^2,3,4^, Sadeed Rashid^11^, Nicklaus Rodriguez^11^, Bushra Samad^2,3,4^, Cole Shaw^3,4^, Austin Sigman^8^, Pratik Sinha^8^, Kevin Tang^11^, Luz Torres Altamirano^11^, Erden Tumurbaatar^10^, Vaibhav Upadhyay^5^, Alyssa Ward^10^, Andrew Willmore^8^, Kristine Wong^7^, Jimmie Ye^14,15,16^, Kimberly Yee^8^, Mingyue Zhou^7^

^1^Division of Hospital Medicine, University of California, San Francisco, CA, USA. ^2^Department of Pathology, University of California, San Francisco, CA, USA. ^3^CoLab, University of California, San Francisco, CA, USA. ^4^Bakar ImmunoX Initiative, University of California, San Francisco, CA, USA. ^5^Department of Medicine, University of California, San Francisco, CA, USA. ^6^Division of Pulmonary and Critical Care Medicine, Department of Medicine, Zuckerberg San Francisco General Hospital and Trauma Center, University of California, San Francisco CA, USA. ^7^Biospecimen Resource Program, University of California, San Francisco, CA, USA. ^8^Division of Pulmonary and Critical Care Medicine, Department of Medicine, University of California San Francisco, San Francisco, California, USA. ^9^Microchemistry, Proteomics and Lipidomics Department, Genentech Inc., 1 DNA Way, South San Francisco, CA, 94080, USA. ^10^Division of Rheumatology, Department of Medicine, University of California, San Francisco, CA 94143, USA. ^11^Helen Diller Family Comprehensive Cancer Center, University of California, San Francisco, CA, USA. ^12^Departments of Otolaryngology and Microbiology & Immunology, Helen Diller Family Comprehensive Cancer Center, CA, USA. ^13^Cardiovascular Research Institute, University of California, San Francisco, San Francisco, CA, USA. ^14^Institute for Human Genetics; Department of Epidemiology and Biostatistics; Institute of Computational Health Sciences; UCSF, University of California, San Francisco, CA, USA. ^15^Parker Institute for Cancer Immunotherapy, University of California, San Francisco, CA, USA. ^16^Chan Zuckerberg Biohub, University of California, San Francisco, CA, USA.

## Methods

### Human subjects

Patients, or a designated surrogate, provided informed consent to participate in the study. The study is approved by the UCSF Institutional Review Board: IRB 20-30497.

### Clinical study design and patient cohort

Clinical study was designed and implemented according to the IMPACC study ((Null et al. 2021)). Patients were recruited from UCSF hospital system and Zuckerberg San Francisco General Hospital and they, or a designated surrogate, provided informed consent to participate in the study. Patients with presumed COVID-19 were enrolled within three days of hospital admission and peripheral blood samples were collected under a protocol approved by the UCSF Institutional Review Board (IRB 20-30497). Patients with confirmed positive SARS-CoV-2 polymerase chain reaction (PCR) were designated as COVID-19 positive cohort (n = 81) and patients without confirmed SARS-CoV-2 PCR were designated COVID-19 negative (n = 7). Healthy donors (n = 11) were recruited (IRB 19-27147) for a single peripheral blood time point and consisted of unexposed patients in a similar age range as the hospitalized cohort. Clinical data and peripheral blood samples were collected at time of enrollment and throughout hospitalization (mainly on days 4, 7, 14, 21, and 28). If escalation of care was required, samples were collected within 24 and 96 hours of care escalation.

### COVID-19 Clinical severity classification

All COVID-19 patients in this study were admitted into the UCSF hospital system and remained there for the duration of our study. By definition, all in-patients reflect a World Health Organization (WHO) COVID-19 severity score of 3 or greater. Patient severity was determined by the clinical team to reflect the WHO COVID-19 severity scoring at each clinical time point throughout in-patient treatment. Based on WHO stratifications (World Health Organization 2021a) and consulting with the treating physician teams, our study combined WHO score 5, 6, and 7 into the most severe clinical group. WHO scores of 3 and 4 correspond to Mild and Moderate groups, respectively.

### Peripheral blood sample collection and processing

Blood samples were collected in one EDTA tube and processed within 6 hours of collection. Whole blood was divided in 540 µL aliquotes then fixed by addition of 756 µL of SmartTube Stabilizer from SmartTube Inc (Fisher Sci. Cat# 501351692). After gentle mixing at room temperature for 10 mins, the samples were transferred to labeled cryovials and immediately carried to −80°C for long term storage.

### Sample Thawing and filtering

Samples were subsequently thawed after being placed 10 min into a 4°C refrigerator then incubated for 15 min in a room temperature water bath. After filtering with 70µm Cell Strainer (Celltreat, Cat# 229483) and washing in 45 ml Milli-Q H2O, samples were counted and barcoded.

### Antibodies and staining procedure

The source for all mass cytometry antibodies can be found in Supplementary Table 1. Antibodies were conjugated to their associated metals with MaxPar X8 labeling reagent kits (Fluidigm) according to manufacturer instructions, diluted with Candor PBS Antibody Stabilization solution (Candor Bioscience, CAT#130 050) supplemented with 0.02% sodium azide, and filtered through an UltrafreeMC 0.1-mm centrifugation filter (Millipore) before storage at 4º C. To reduce tube-to-tube pipetting variations, part of the signaling antibody panel came from lyophilized antibody cocktail, made at Stanford University as previously described ((Han et al. 2018)). Surface and intracellular master antibody cocktails were made and kept at -80º C in order to stain up to 600 samples.

### Mass-tag cellular barcoding

Prior to antibody staining, mass tag cellular barcoding of prepared samples was performed by incubating cells with distinct combinations of isotopically-purified palladium ions chelated by isothiocyanobenzyl-EDTA as previously described ((Zunder et al. 2015)). After counting, 1*10^6^ cells from aliquot were barcoded with distinct combinations of stable Pd isotopes for 15 min at room temperature on a shaker in Maxpar Barcode Perm Buffer (Fluidigm, cat#201057). Cells were washed twice with cell staining media (PBS with 0.5% BSA and 0.02% NaN3), and pooled into a single 15 ml tube.

### Mass cytometry staining

Barcoded cells were stained with Fc Receptor Blocking Solution (BioLegend, Cat#422302) at 20 mg/ml for 5 min at RT on a shaker. Surface antibody cocktail is then added with a 500 ul final reaction volume for 30 min at RT on a shaker. Following staining, cells were washed twice with cell staining media. Before intracellular staining, cells were permeabilized for 10 min with methanol at 4ºC. Methanol is then removed by washing the cells 2 times with cells staining media. Intracellular cocktail is then added to the cells in 500 uL final reaction volume for 1 hour at RT on a shaker. Cells were washed twice in cell staining media to remove antibodies excess and then stained with 1mL of 1:4000 191/193Ir Iridium intercalator solution (Fluidigm, Cat#201192B) diluted in PBS with 4% PFA overnight. Before mass cytometry run, cells were washed once with cell staining media, and twice with Cell Acquisition Solution (Fluidigm, Cat# 201240).

### Mass Cytometry

Mass cytometry samples were diluted in Cell Acquisition Solution containing bead standards (Fluidigm, Cat#201078) to approximately 10^6^ cells/mL and then analyzed on a Helios mass cytometer (Fluidigm) equilibrated with Cell Acquisition Solution. Approximately 0.5×106 cell events were collected for each sample at an even rate of 400–500 events/second.

### Data normalization and de-barcoding

Bead standard data normalization and de-barcoding of the pooled samples into their respective conditions was performed using the R package from the PICI institute available at https://github.com/ParkerICI/premessa.

### Quality control inclusion and exclusion criteria

In order to ensure high quality sample collection, processing, and staining across the cohort we developed a set of inclusion criteria required for each sample to be used in our data analysis. We processed and ran CyTOF on 498 peripheral blood samples. After debarcoding and normalization, samples were uploaded to Cell Engine to assess adequate staining and cell number. Each barcode plate was run with a healthy PB control sample aliquoted from two healthy donors to validate staining and for normalization between barcode plates. If the control PB sample failed to stain the major immune cell populations (T cell, B cell, granulocytes, monocytes), no samples from that barcode plate were included. Individual samples were then assessed for CD45+ composition (>50% CD45+ staining required), cell abundance (>5,000 cells per sample required), and representation of the major immune cell populations (T cell, B cell, granulocytes, monocytes). 230 samples passed QC and were used in the batch normalization.

## QUANTIFICATION AND STATISTICAL ANALYSIS

### Batch normalization

All manually gated immune cells (CD45+) from samples meeting our inclusion criteria (n = 230) were downloaded as FCS files from cellEngine. Premessa (Gherardini 2021) (https://github.com/ParkerICI/premessa) and cytofCore (R. Bruggner and M. Linderman and R. Finck 2021) (https://github.com/nolanlab/cytofCore) were used to harmonize panels between runs, and CytoNorm (Van Gassen 2021) (https://github.com/saeyslab/CytoNorm) were utilized to correct for batch effect. All markers were used for batch effect normalization, except for Ki-67, which failed for several CyTOF runs and were excluded in the final data. Samples were separately normalized to control 1 and 2, and subsequently combined into one final data set of normalized FCS files.

### Manual gating

Batch effect normalized FCS files were uploaded to Cell Engine for manual gating. Major immune cell populations were identified based on prior gating strategy (Allen et al. 2020). T cell subsets were further identified based on phenotypic markers specified in prior publication that suggested these specific subtypes could play a role in COVID-19 severity (Mathew et al. 2020).

### t-SNE visualization

The multiparameter dimensionality reduction method t-distributed stochastic neighbor embedding (t-SNE) was employed to visualize major shifts in immune distribution between COVID-19 positive, COVID-19 negative, and healthy individuals. CD45+ immune cells from healthy peripheral blood samples were compared to day 0 (D0) peripheral blood samples from COVID-9 positive and negative individuals and respective groups were concatenated into a single FSC file which was then used in the t-SNE algorithm on Cell Engine (cellengine.com). Only phenotypic markers were used as analysis channels and no phospho-signaling channels were input into the t-SNE visualization. The default settings for t-SNE plot were utilized and a default of 90 nearest neighbors (k) was used. Manually gated immune cell populations were used to color the t-SNE plot to identify representative immune populations on the plot.

### Defining groups and samples

For intra-patient resolution analyses, we defined three different groups; patients who were discharged within 30 days of enrollment in the study (<30 days), patients who were discharged after 30 days of enrollment in the study (>30 days), and patients who died. For patients who were discharged <30 days, the last sample (tp2) had to be obtained within 7 days of discharge. For patients who were discharged >30 days and patients who died, the last sample (tp2) had to be obtained within 50 days of discharge. For all groups, the first sample (tp1) had to be obtained within 14 days of enrollment. For intra-patient ventilation recovery analysis, samples had to be obtained within 7 days of the point of interest, e.g. going on a ventilator / coming off a ventilator. For all comparisons; if multiple samples fulfilled the requirements, we used the sample closest to the event of interest. The number of patients and specific sampling timepoints used for each analysis are illustrated in the supplementary figures.

### Statistical analysis

All statistical tests were performed in R (Team and Others 2013; RStudio Team 2016). The non-parametric Wilcoxon rank sum test was utilized to compare immune population frequencies, median protein expression values, and median signaling molecule values between groups of interest. For intra-patient analysis, we used the paired Wilcoxon rank sum test. For multiple testing corrections, we applied Benjamini-Hochberg correction and statistical differences were declared significant at FDR < 0.1. When multiple testing was not applied, statistical differences were declared significant at P < 0.05. Most of the plots were produced with the R package ggplot2 (Wickham 2016).

## Supplemental Tables

**Supplemental table 1:** Patient demographics and clinical parameters, e.g.

WHO score at time of sampling, max WHO score, hospital length of stay, ventilation duration, etc. (patient_demographics.xlsx)

**Supplemental table 2:** Summary of patient demographics

(Patient_demographics_summary.xlsx)

**Supplemental table 3:** Healthy individuals demographics (healthy_demographics.xlsx)

**Supplemental table 4:** Antibody panel (antibody_panel.xlsx)

---

**Supplemental figure 1.**
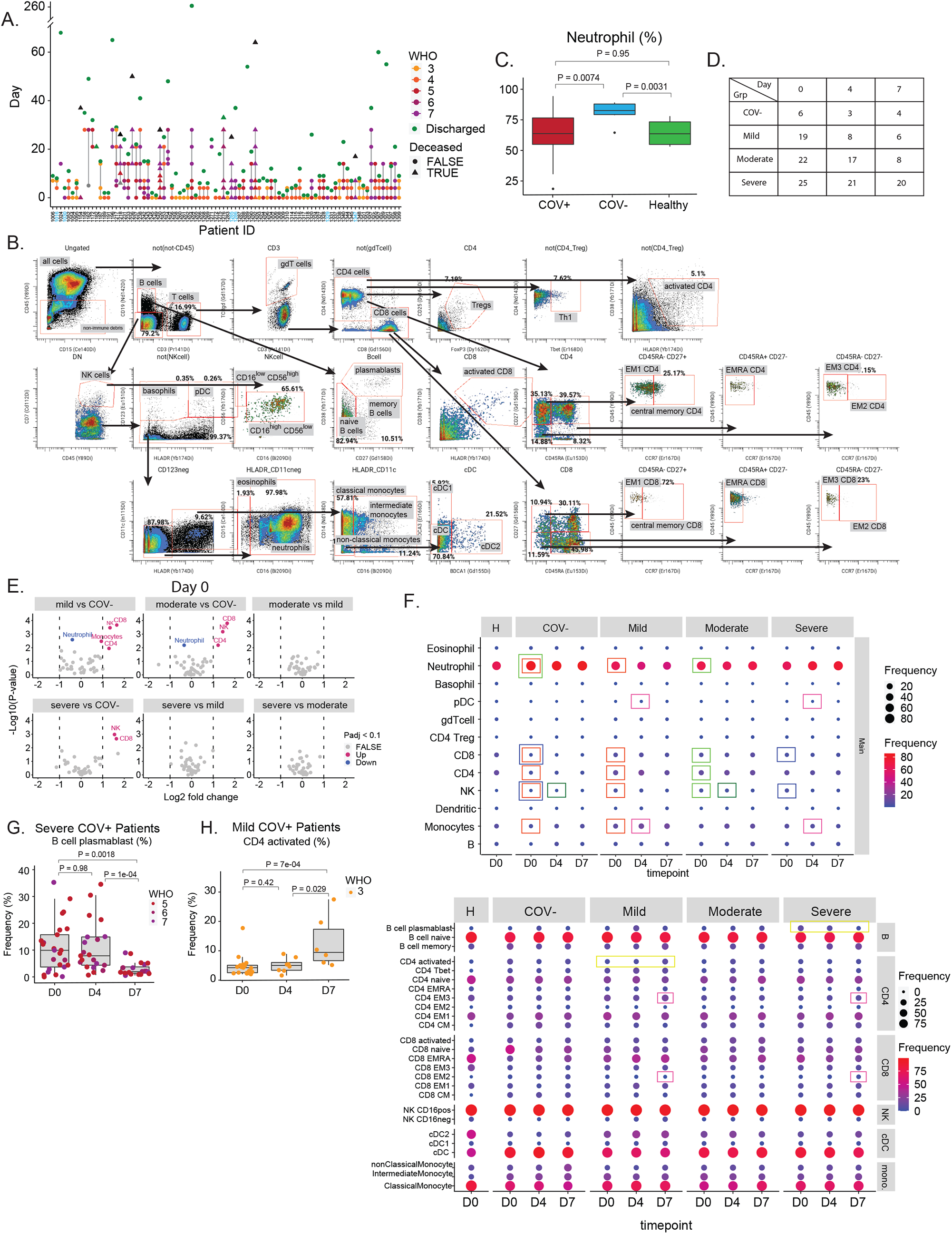
**A)** All patient samples included in study (220 samples from 89 patients). COVID-19 patients shown in black and COVID-19 negative patients in blue. Points indicate sample timepoint and are coloured according to WHO score. Green points indicate the day of discharge, while triangles indicate patients that died. **B)** Gating strategy for manual immune cell gates. **C)** Frequency of neutrophils in COVID-19 positive (COV+), COVID-19 negative (COV-) patients, and healthy controls at D0. P-values obtained by Wilcoxon Rank Sum Test. **D)** Number of samples at each time point (D0, D4, and D7) for each COVID-19 severity group and for COVID-19 negative patients. **E)** Differential expression analysis of immune cell populations between COVID-19 severity groups and COVID-19 negative patients at D0. The log2 fold changes are plotted against the negative log10(p-values). P-values obtained by Wilcoxon Rank Sum Test. Colors indicate if cell populations are significantly down- (blue) or upregulated (purple) or not differentially expressed (FALSE, grey) after Benjamini-Hochberg correction, FDR < 0.1. **F)** Immune cell population abundance at D0, D4, and D7 in COVID-19 patients divided into severity groups based on their WHO score, as well as in COVID-19 negative patients, and healthy individuals at D0. P-values obtained by Wilcoxon Rank Sum Test, followed by Benjami-ni-Hochberg correction with FDR < 0.1. Immune cell populations that are significantly different after BH correction (across time points within groups or cross-sectional at the same time point between groups) are highlighted with coloured boxes corresponding to the time point and group of comparison. All comparisons between patients and healthy individuals at D0 are illustrated with p-values in main figure 1C. **G+H)** Frequency of B cell plasmablasts (G) and CD4 activated T cells (H) in patients suffering from severe and mild disease, respectively. P-values obtained by Wilcoxon Rank Sum Test.

**Supplemental figure 2.**
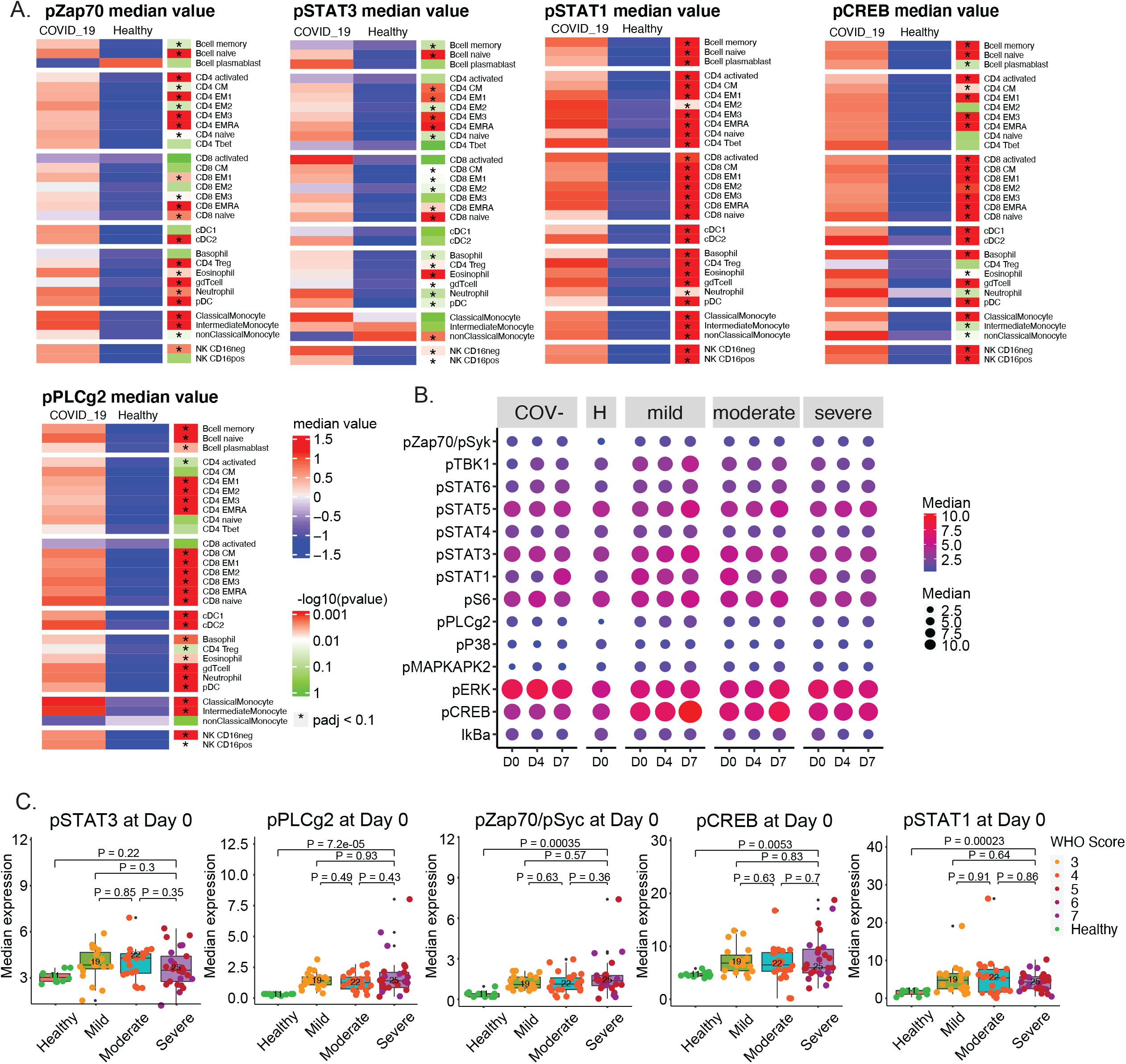
**A)** Expression of significant changing signaling molecules from 2B in all CD45+ cell population subsets at D0 in COVID-19 patients and healthy individuals. Median expression values have been centered on heatmap. P-values obtained by Wilcoxon Rank Sum Test, followed by Benjamini-Hochberg correction with FDR < 0.1. **B)** Median signaling molecule values at D0, D4, and D7 in COVID-19 patients divided into severity groups based on their WHO score, as well as in COVID-19 negative patients, and healthy individuals at D0. **C)** Specific comparisons from S1B. P-values obtained by Wilcoxon Rank Sum Test.

**Supplemental Figure 3.**
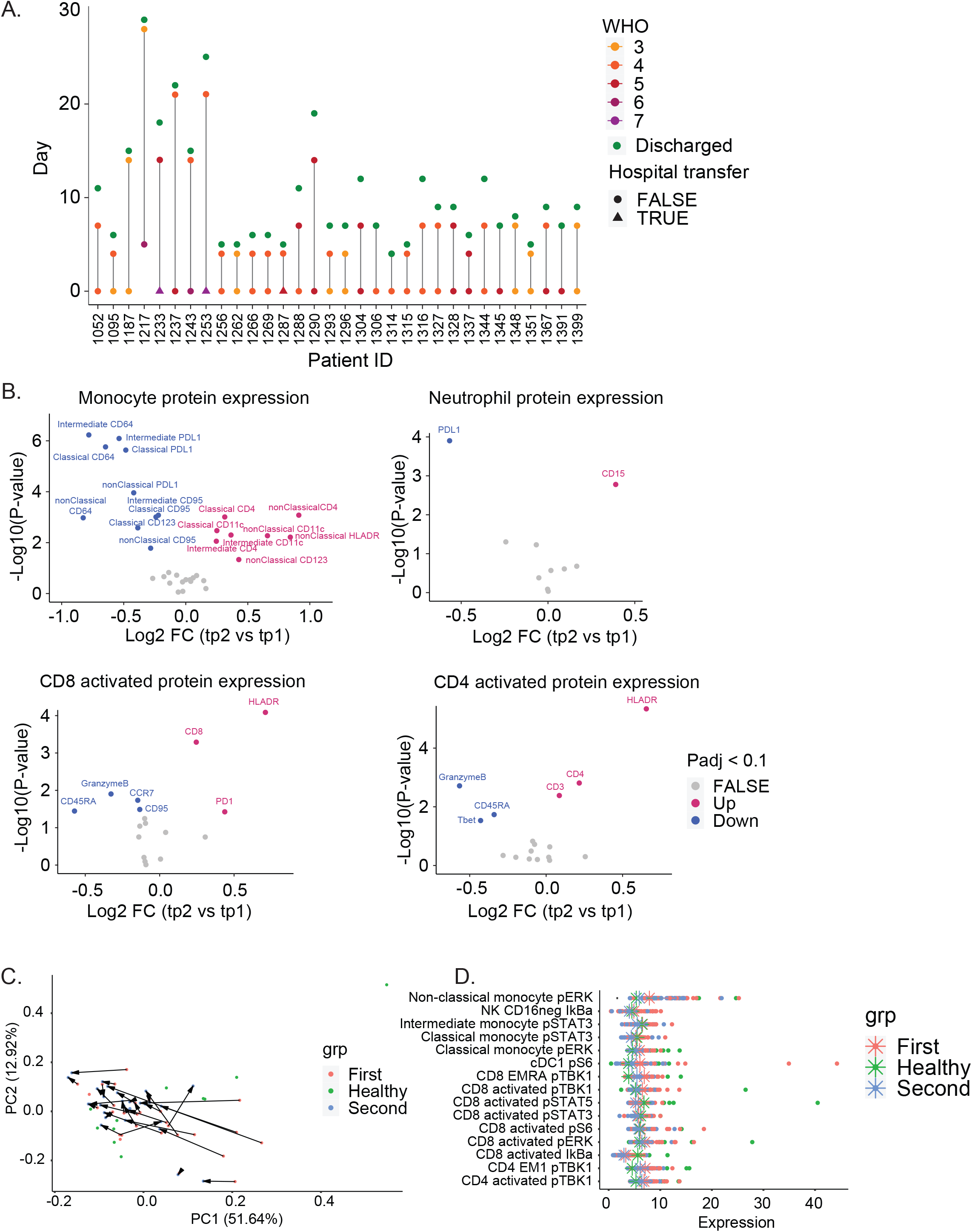
**A)** Samples used for intra-patient analysis in Figure 3 of patients that are discharged from the hospital within 30 days of admission (n = 32). Points indicate sample timepoint and are coloured according to WHO score. Green points indicate the day of discharge. **B)** Paired differential expression analysis of protein expression on monocyte subsets, neutrophil, CD8- and CD4 activated T cells between the first (tp1) and second (tp2) timepoints illustrated in 3A (paired Wilcoxon Rank Sum Test). The log2 fold changes (tp2 vs tp1) are plotted against the negative log10(p-values). Colors indicate if cell populations are significantly down- (blue) or upregulated (purple) from tp1 to tp2 or not differentially expressed (FALSE, grey) after Benjamini-Hochberg correction, FDR < 0.1. **C)** Principal component analysis of significant signaling molecules in 3I for tp1, tp2, and healthy controls. **D)** Expression of signaling molecules (from 3I and S3C) for tp1, tp2, and healthy controls. Stars indicate median value for each group.

**Supplemental Figure 4.**
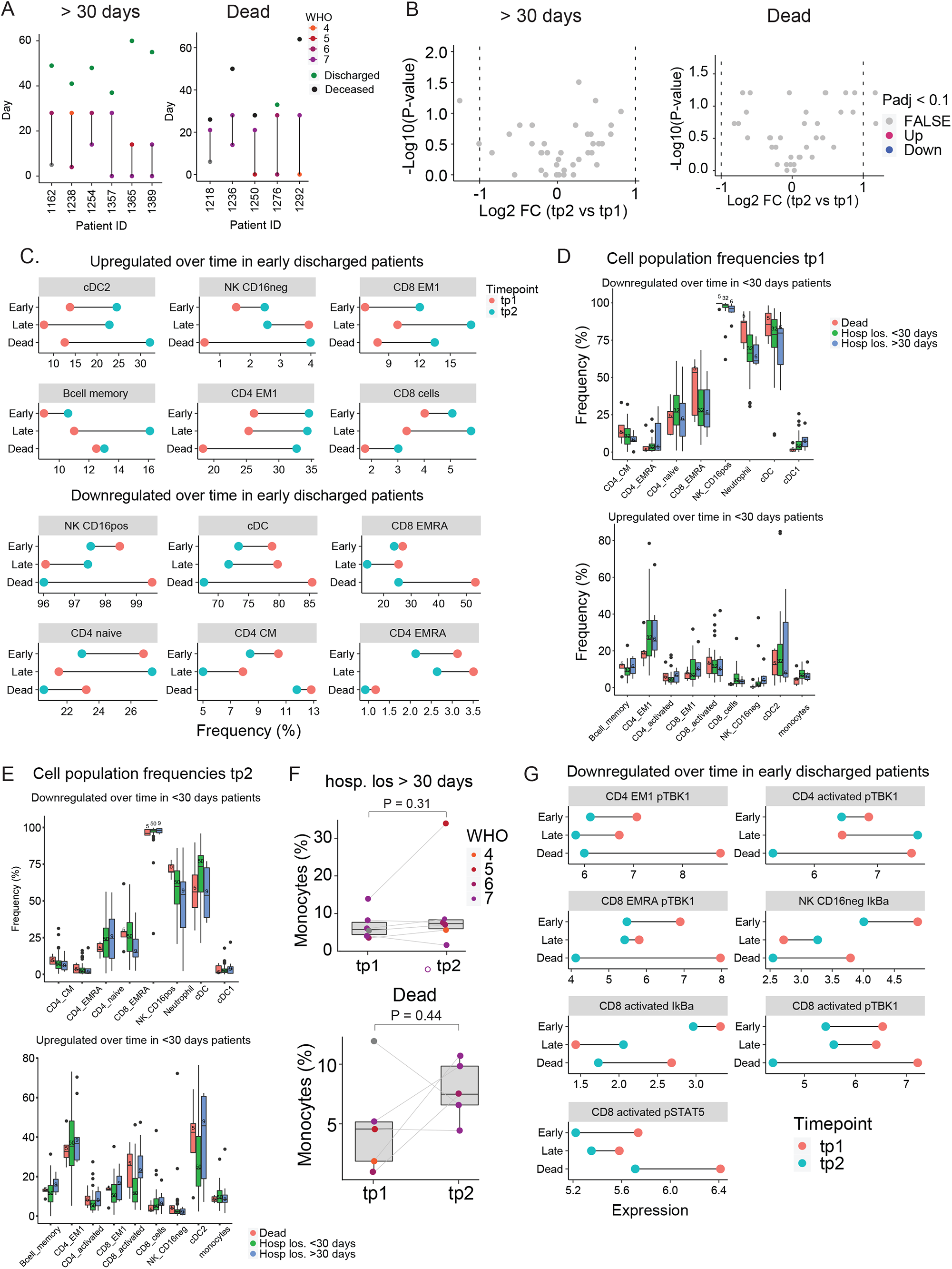
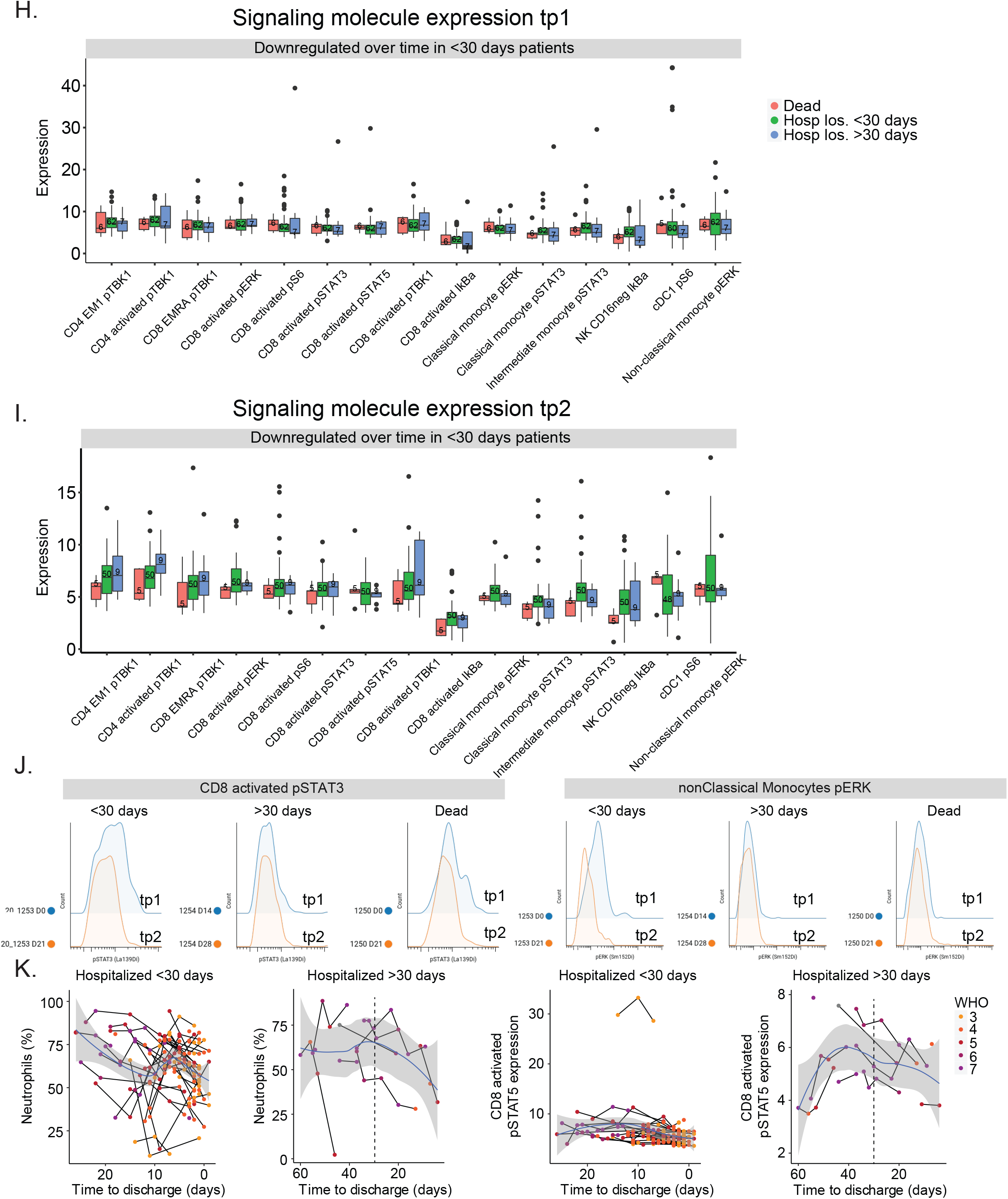
**A)** Samples used for intra-patient analysis in Figure 4 of patients that are discharged after 30 days (left, n = 6) and patients that die (right, n = 5). Points indicate sample timepoint and are coloured according to WHO score. Green points indicate the day of discharge. **B)** Paired differential expression analysis of immune cell populations between the first (tp1) and second (tp2) timepoints illustrated in 4A (paired Wilcoxon Rank Sum Test). The log2 fold changes (tp2 vs tp1) are plotted against the negative log10(p-values). Colors indicate if cell populations are significantly down- (blue) or upregulated (purple) from tp1 to tp2 or not differentially expressed (FALSE, grey) after Benjamini-Hochberg correction, FDR < 0.1. **C)** Median cell population frequencies at tp1 (red) and tp2 (blue) for patients that are discharged <30 days, >30 days, and deceased. **D+E)** Cell population frequencies at tp1 (D) and tp2 (E) for patients that are discharged <30 days, >30 days, and deceased. **F)** Frequency of monocytes at tp1 and tp2 for patients that are discharged after 30 days or die. Lines connect samples from the same patient. P-values obtained by paired Wilcoxon Rank Sum Test. **G)** Median expression of signaling molecules at tp1 (red) and tp2 (blue) for patients that are discharged <30 days, >30 days, and deceased. **H+I)** Expression of signaling molecules at tp1 (H) and tp2 (I) for patients that are discharged <30 days, >30 days, and deceased. **J)** Expression of pSTAT3 in activated CD8 T cells (left) and pERK in non-classical monocytes (right) at tp1 (blue) and tp2 (orange) for representative patients that are discharged <30 days, >30 days, and deceased. **K)** Neutrophil frequencies (left plots) and CD8 activated pSTAT5 expressions (right plots) relative to time to discharge in all samples from patients who are discharged <30 days (n = 142 samples) or >30 days (n = 30 samples). Black lines connect samples from the same patient. Blue lines and grey shadow represent the best fitted smooth line and 95% confidence interval.

**Supplemental Figure 5:**
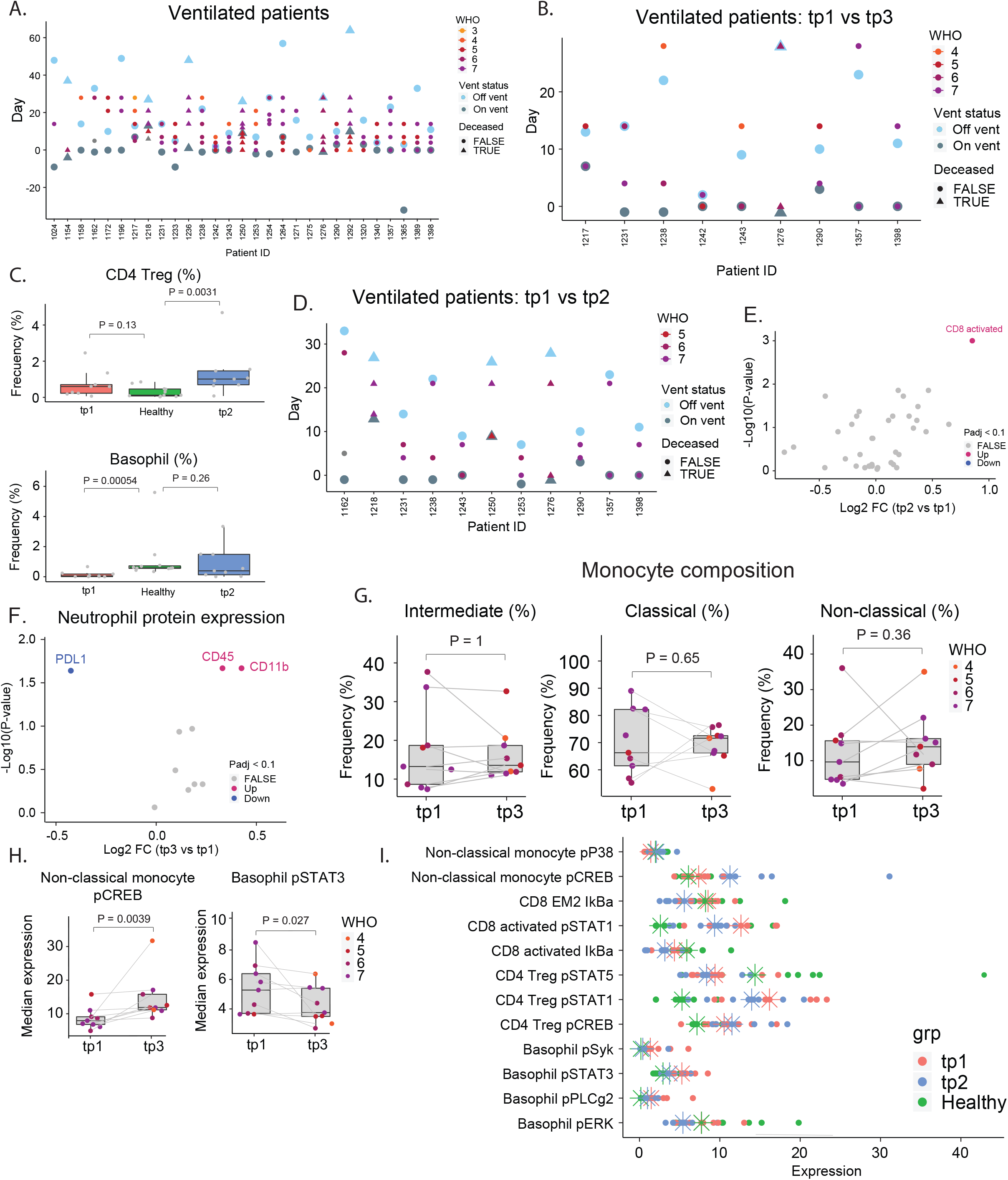
**A)** All samples for patients that have been put on a ventilator. Dark blue points indicate when a patient is put on a ventilator. Light blue points indicate when a patient is taken off a ventilator. **B)** Samples used for intra-patient analysis between tp1 and tp3 in Figure 5 of patients that have been put on a ventilator (n = 9). Points indicate sample timepoint and are coloured according to WHO score. Dark blue points indicate when a patient is put on a ventilator. Light blue points indicate when a patient is taken off a ventilator. **C)** Frequencies of CD4 Tregs and Basophils at tp1, tp3, and in healthy controls. P-values obtained by Wilcoxon Rank Sum Test. **D)** Samples used for intra-patient analysis between tp1 and tp2 in Figure 5 for patients that have been put on a ventilator (n = 11). Points indicate sample timepoint and are coloured according to WHO score. Dark blue points indicate when a patient is put on a ventilator. Light blue points indicate when a patient is taken off a ventilator. **E)** Paired differential expression analysis of immune cell populations between the first (tp1) and second (tp2) timepoints illustrated in 5A (paired Wilcoxon Rank Sum Test). The log2 fold changes (tp2 vs tp1) are plotted against the negative log10(p-values). Colors indicate if cell populations are significantly down- (blue) or upregulated (purple) from tp1 to tp2 or not differentially expressed (FALSE, grey) after Benjamini-Hochberg correction, FDR < 0.1. **F)** Paired differential expression analysis of protein expression on neutrophils between the first (tp1) and third (tp3) timepoints illustrated in 5A (paired Wilcoxon Rank Sum Test). The log2 fold changes (tp3 vs tp1) are plotted against the negative log10(p-values). Colors indicate if cell populations are significantly down- (blue) or upregulated (purple) from tp1 to tp3 or not differentially expressed (FALSE, grey) after Benjamini-Hochberg correction, FDR < 0.1. **G)** Frequencies of monocyte subsets at tp1 and tp3. P-values obtained by paired Wilcoxon Rank Sum Test. **H)** Expression of pSTAT3 in Basophils and pCREB in non-classical monocytes at tp1 and tp3. Lines connect samples from the same patient. P-values obtained by paired Wilcoxon Rank Sum Test. **I)** Expression of signaling molecules in 5M for tp1, tp2, and healthy controls. Stars indicate median value for each group.

**Supplemental Figure 6:**
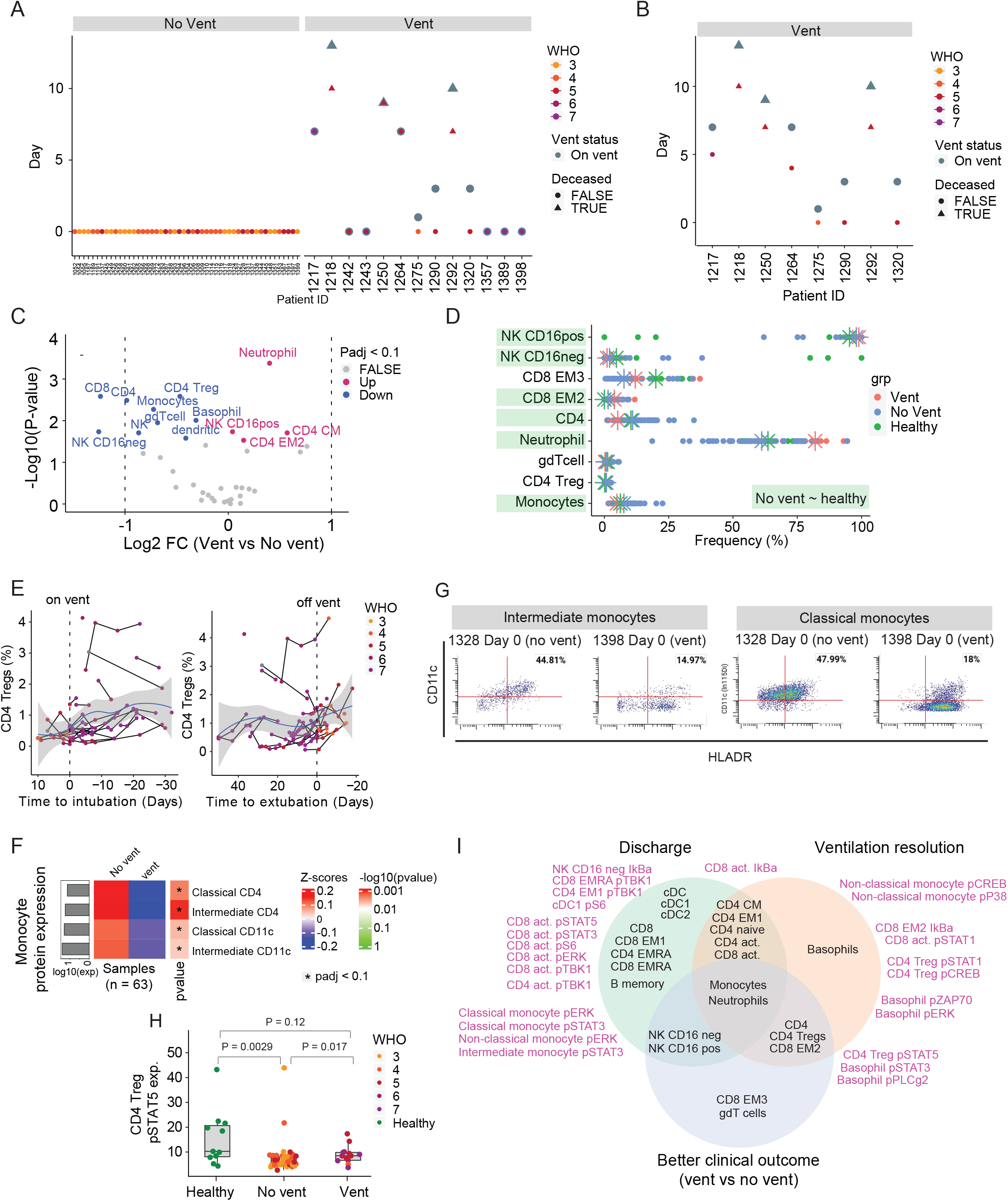
**A)** Samples used for inter-patient analysis in Figure 6. For ventilated patients (n = 13), the latest sample before the patient is put on a ventilator or, if available, the sample at the day of ventilation is used. For non-ventilated patients (n = 50), D0 is used. **B)** Samples obtained prior to ventilation (n = 8). **C)** Differential expression analysis of immune cell populations between ventilated (from S6B) and non-ventilated patients (from S6A) (Wilcoxon Rank Sum Test). The log2 fold changes (vent vs no vent) are plotted against the negative log10(p-values). Colors indicate if cell populations are significantly down- (blue) or upregulated (purple) for vent vs no vent or not differentially expressed (FALSE, grey) after Benjamini-Hochberg correction, FDR < 0.1. **D)** Population frequencies of significant immune cell subsets in 6B for ventilated-, non-ventilated patients, and healthy controls. Stars indicate median value for each group. Cell populations are highlighted in green if non-ventilated patients are closer to healthy controls than ventilated patients. **E)** CD4 Treg frequencies relative to intubation / extubation in all samples from ventilated patients. Black lines connect samples from the same patient. Blue lines and grey shadows represent the best fitted smooth line and 95% confidence interval. Dotted lines intersect the x-axis at day of intubation / extubation. **F)** Protein expression on monocyte subsets in ventilated- and non-ventilated patients. Mean protein expression values have been log10 transformed, scaled, and centered on heatmap. Bars indicate mean protein expression across all samples. Only significant proteins are shown (Wilcoxon Rank Sum Test, Benjamini-Hochberg correction with FDR < 0.1). **G)** Scatter plots of CD11c and HLA-DR expression on intermediate (left) and classical monocytes (right) in representative patients. **H)** Expression of pSTAT5 in CD4 Tregs in non-ventilated and ventilated patients as well as healthy individuals. P-values obtained by Wilcoxon Rank Sum Test. **I)** Significantly changing immune cell populations (black text) and signaling molecules (purple) accompanying discharge (green), ventilation resolution (orange), and better clinical outcome (blue).

